# A Novel Hydrogen Polysulfide Donor Alleviates Neuropathic Pain by Enhancing A-Type Potassium Currents via LIMK1-Cofilin Mediated Interaction between Filamin A and Kv4.2

**DOI:** 10.64898/2026.05.25.727751

**Authors:** Ting Kang, Yan Jiao, Shirong Lai, Shouwei Tao, Chunxia Chen, Zhihua Cao, Fei Yan, Yanwen Ding, Xuehan Li, Bowen Ke

## Abstract

Hydrogen persulfide (H_2_S_2_) is an important endogenous signaling molecule, holding significant therapeutic potential across diverse disease models due to its potent antioxidant and redox-regulating properties. Herein, we report the synthesis, characterization, and in vivo evaluation of an esterase responsive H_2_S_2_ donor, **HPD1**. It reduced mechanical and cold allodynia at 14 mg/kg (i.p.) in chronic constriction injury and paclitaxel-induced neuropathic pain models. Moreover, **HPD1** exhibited negligible systemic toxicity and behavioral side effects even at 28 mg/kg. Electrophysiological tests showed that **HPD1** suppressed PTX-induced hyperexcitability in dorsal root ganglion (DRG) neurons by specifically potentiating A-type potassium currents (*I*_A_). Mechanistically, we demonstrate that **HPD1** activates LIMK1, which inactivates cofilin and stabilizes F-actin, thereby promoting the interaction between the actin-binding protein Filamin A and Kv4.2. Furthermore, both the **HPD1**-induced increase in Filamin A-Kv4.2 co-localization and the subsequent restoration of *I*_A_ density in DRG neurons, as well as the analgesic effect of **HPD1** were dampened by pharmacological inhibition of LIMK1 with BMS-5. This work has developed a new generation of H_2_S_2_ donors, demonstrated the analgesic efficacy of **HPD1**, and uncovered the novel LIMK1-cofilin-Filamin A-Kv4.2 dependent mechanism that restores *I*_A_, thus providing a reliable therapeutic strategy for neuropathic pain based on H_2_S_2_ donors.

## 1. Introduction

Hydrogen persulfide (H_2_S_2_) is a reactive sulfur species that has recently gained recognition as an endogenous signaling molecule with distinct biological properties that set it apart from its more extensively studied counterpart, hydrogen sulfide (H_2_S)^1^. As a member of the reactive sulfur species family, H_2_S_2_ contains a sulfur-sulfur bond, which imparts higher electrophilicity and potent antioxidant and redox-modulating activities^2^. Emerging evidence suggests that H_2_S_2_ plays critical roles in regulating cellular redox homeostasis, protein persulfidation, and ion channel function, implicating it in various physiological and pathological processes^1,3^. However, the therapeutic potential of H_2_S_2_ has long been hindered by its inherent instability and the limited availability of donors with controlled release kinetics^4,5^. Consequently, the development of well-defined H_2_S_2_ donors is of great interest for both biological investigations and therapeutic development.

Our previous finding that an esterase-sensitive H_2_S_2_ donor exerts analgesic effects in acute pain models^6^ prompted us to examine whether H_2_S_2_ donors might also alleviate neuropathic pain, which affects approximately 7-10% of the global population. Neuropathic pain, resulting from lesions or diseases of the somatosensory nervous system, is often inadequately managed by current therapeutics due to limited efficacy and dose-limiting side effects^7,8^. Hyperexcitability of primary afferent neurons is frequently observed in various animal models of neuropathic pain, and this pathological state is typically accompanied by a diminished K^+^ current^9–12^. Potassium currents serve as the predominant outward conductance in neurons, promoting membrane hyperpolarization and thereby reducing neuronal excitability^13^. Based on their kinetic properties and pharmacological profiles, K^+^ currents are generally classified into at least two distinct categories: the sustained delayed-rectifier potassium current (*I*_K_) and the transient A-type potassium current (*I*_A_). Among these, *I*_A_ plays a critical role in determining action potential threshold, waveform, and firing frequency^14^. In mammals, *I*_A_ is mediated by voltage-gated potassium channel (Kv) α-subunits encoded by the genes Kv1.4, Kv3.4, Kv4.1, Kv4.2, and Kv4.3. Previous studies have shown that DRG neurons express several of these Kv α-subunits, and that peripheral nerve injury leads to their downregulation^11^. The function and surface expression of these channels are dynamically regulated by their interaction with auxiliary subunits, including cytosolic Kv channel-interacting proteins (KChIPs) and dipeptidyl peptidase-related proteins (DPPs), as well as with cytoskeletal proteins such as the actin-binding protein Filamin A^15–17^. Nevertheless, the therapeutic potential of targeting these modulatory interactions, either directly or indirectly, to restore *I*_A_ for pain relief has yet to be explored.

LIM-domain containing protein kinases (LIMK1 and LIMK2) regulate actin dynamics by phosphorylating and inactivating cofilin, an actin-severing protein, thereby stabilizing filamentous actin (F-actin)^18^. This cytoskeletal remodeling plays a critical role in synaptic plasticity, neuronal morphology, and the trafficking of membrane proteins, including ion channels and receptors^19,20^. The LIMK1-cofilin pathway plays a complex, context-dependent role in pain processing. While some studies have reported that inhibition of LIMK1 attenuates inflammatory and neuropathic pain^21,22^, others suggest that appropriate levels of LIMK1 activity are required for maintaining normal neuronal function^23^. This apparent discrepancy may arise from differences in pain models, cell types, or the temporal dynamics of LIMK1 activation. Regardless, the downstream mechanisms by which LIMK1-cofilin signaling modulates ion channel function in DRG neurons remain poorly defined.

Here, we present the synthesis, release kinetics, and systematic safety evaluation of a novel esterase-triggered H_2_S_2_ prodrug, **HPD1**, and demonstrate its analgesic efficacy and mechanism of action in neuropathic pain models. In paclitaxel (PTX)-induced and chronic constriction injury (CCI) models, **HPD1** effectively attenuated mechanical allodynia and cold hypersensitivity without affecting locomotor activity or affective behaviors. Whole-cell patch-clamp recordings revealed that **HPD1** selectively normalized hyperexcitability of DRG neurons by restoring the pathologically suppressed *I*_A_, without altering sodium or calcium currents. Mechanistically, we demonstrate that **HPD1** activates the LIMK1-cofilin pathway, leading to phosphorylation of LIMK1 (Thr508) and cofilin (Ser3), and thereby stabilizes F-actin. This cytoskeletal remodeling enhances the physical interaction between the actin-binding protein Filamin A and Kv4.2. The functional relevance of this pathway was validated by the selective LIMK1/2 inhibitor BMS-5, which blocked **HPD1**-induced LIMK1/cofilin phosphorylation, Filamin A-Kv4.2 interaction, *I*_A_ potentiation, and analgesic effects. Collectively, our findings identify **HPD1** as a safe and effective analgesic that alleviates neuropathic pain by restoring *I*_A_ through a previously unrecognized LIMK1-cofilin-Filamin A-Kv4.2 signaling axis, establishing H_2_S_2_ donors as a promising therapeutic strategy for neuropathic pain.

## 2. Results

### 2.1 A Generic Donor Scaffold for Esterase-Triggered H_2_S_2_ Release

In developing our new generation H_2_S_2_ donors, we focused on designs incorporating “carrier” moieties with known safety profiles into the donor scaffold. Employing a design strategy similar to that of previously reported esterase-sensitive persulfide donors^24^, we exploited the bis(2-hydroxyethyl) disulfide as a key intermediate. An ester group was introduced to mask the hydroxy group, generating a stable precursor. As illustrated in **Scheme 1**, the key to this design is an esterase-labile trigger group, which sets off the following decomposition cascade. Upon enzymatic hydrolysis, these molecules undergo intramolecular elimination, releasing two equivalents each of an acetaldehyde (CH_3_CHO) and a carboxylic acid (R–COOH) that include endogenous metabolites, food-grade additives, and pharmaceutical compounds (e.g., acetic acid, phenylacetic acid, acetylsalicylic acid). This design thus offers a controlled release mechanism for H_2_S_2_ under physiological conditions, with the released byproducts posing no safety risks.

**Scheme 1.**
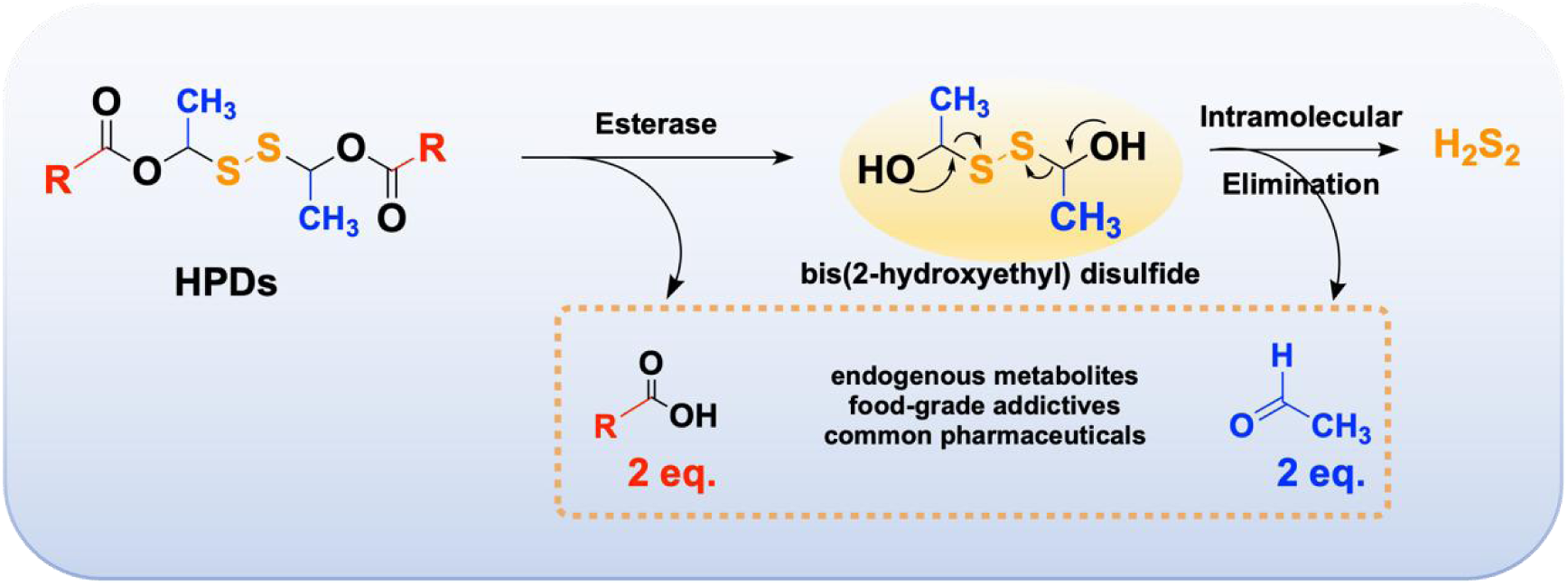
Design and Enzyme-Triggered Release Mechanism of the H_2_S_2_ Donor Scaffold.

Guided by this strategy, we synthesized the donor molecules **HPD1**-**HPD3** via a concise two-step sequence (**Scheme 2**). The halide precursor **a1** (for **HPD1**) was obtained commercially, whereas precursors **a2** and **a3** (for **HPD2** and **HPD3**) were synthesized from readily available starting materials (**Scheme S1**). The general synthetic route involved nucleophilic substitution of the halide precursors (**a1**-**a3**) with potassium thioacetate (KSAc) in acetone at room temperature to afford the corresponding thioacetate intermediates (**b1**-**b3**), followed by a pivotal iodine-catalyzed oxidative homocoupling in DCM/DMF at −10 °C to construct the disulfide bond, delivering the target H₂S₂ donors (**HPD1**-**HPD3**).

**Scheme 2.**
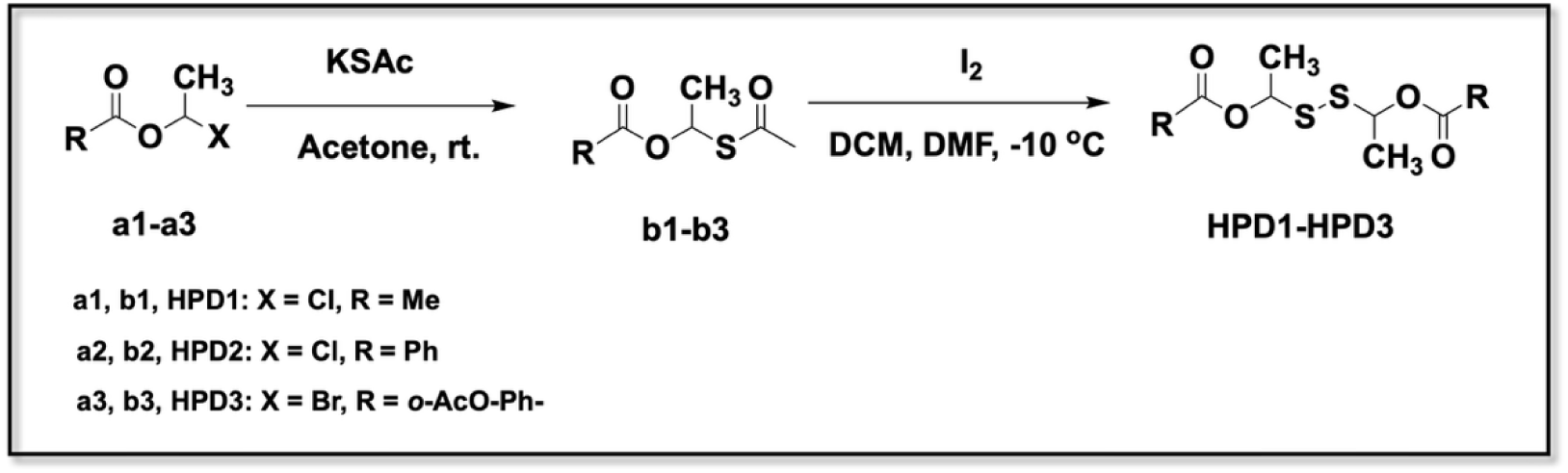
Synthetic Routes of H_2_S_2_ donors **HDP1-HPD3**. DCM = dichloromethane, DMF = N, N-dimethylformamide, KSAc = potassium thioacetate.

### 2.2 Identification and Characterization of H_2_S_2_ Donors

To identify the optimal donor for further study, we compared the solubility or release characteristics of the three candidates. Among them, both **HPD1** and **HPD3** exhibited substantially improved solubility compared to our previously reported donor **HPS-2** (0.0026 mg/mL)^6^, with approximately 700-fold and 7-fold enhancements, respectively. The solubility data for all donors are summarized in **Table S1**. Since release capacity is a critical determinant of pharmacological efficacy for a prodrug, we next performed a semi-quantitative assessment of H_2_S_2_ release from all three candidates using the fluorescent probe DSP-3^4^. Under identical conditions (1 U PLE, 37 °C, 15 min), released H_2_S_2_ was captured by the probe and quantified by fluorescence intensity, with signals normalized to the corresponding vehicle-treated controls. As shown in **Fig. 1A**, the relative fluorescence intensities were 15.93 ± 0.80 for **HPD1**, 0.59 ± 0.10 for **HPD2** and 4.13 ± 0.37 for **HPD3**, with **HPD1** yielding values comparable to the positive control Na_2_S_2_ (17.88 ± 0.62). Notably, in the absence of PLE, none of the donors produced significant fluorescence (**Fig. S1A**), ruling out appreciable background signals from the donors alone. Therefore, **HPD1** was chosen for further studies owing to its superior solubility and release efficiency.

**Figure 1.**
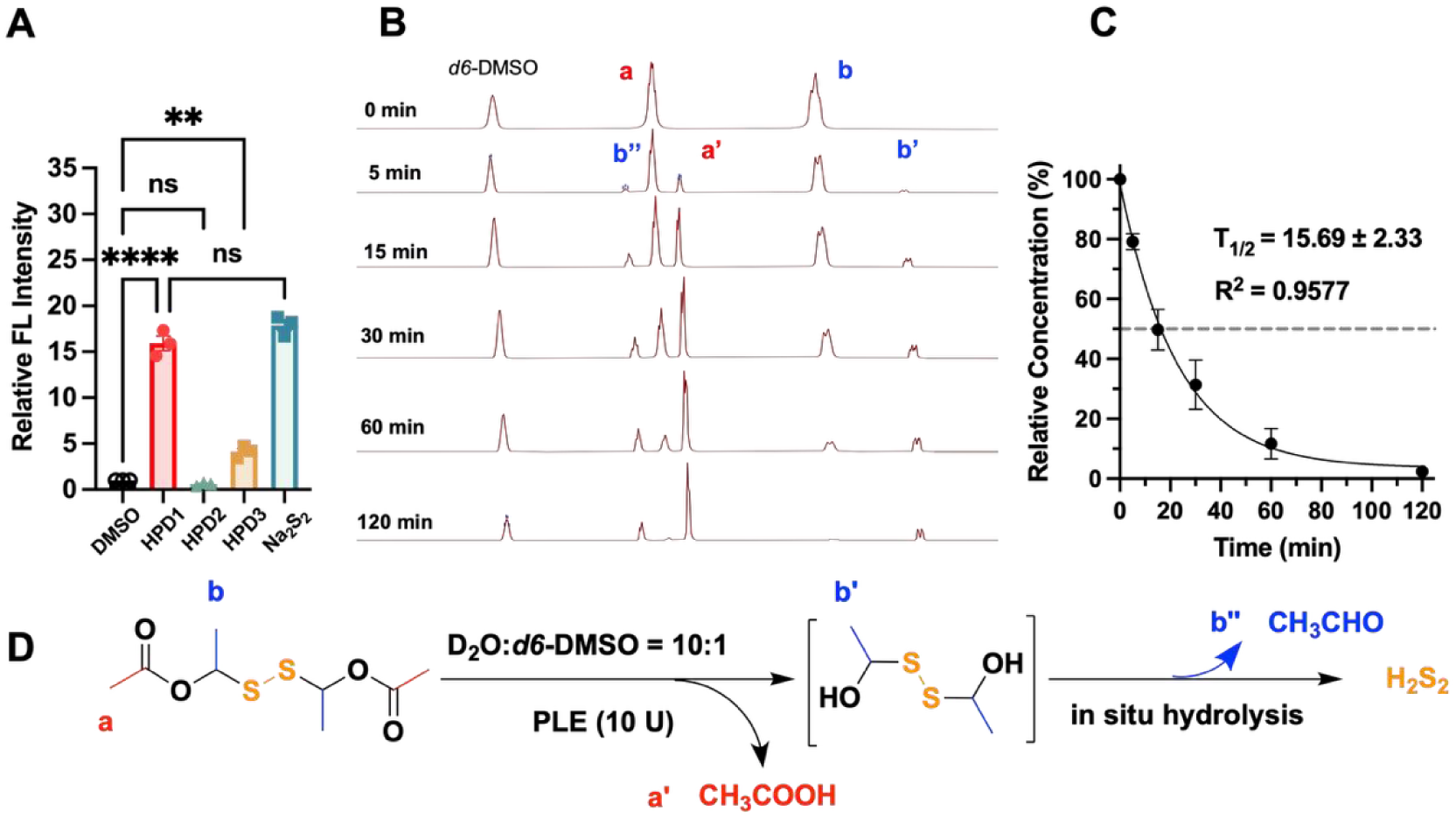
Fluorescence-based screening of HPDs and characterization of HPD1 release kinetics and mechanism. (A) Semi-quantitative fluorescence analysis of H_2_S_2_ release from HPDs in the presence of PLE, with Na_2_S_2_ as positive control, using DSP-3 probe λ_ex_/λ_em_ = 490/515 nm. (B) ^1^H NMR spectra (400 MHz, D₂O/*d_6_*-DMSO = 10:1) tracking the time-dependent decomposition of **HPD1** (10 mM) upon incubation with PLE (10 U). (C) Pseudo-first-order kinetic analysis of **HPD1** decomposition and corresponding half-life (T_1/2_) calculated from ^1^H NMR monitoring. (D) Proposed mechanism for H_2_S_2_ release from **HPD1**. Data are mean ± SEM (n = 3); ns, not significant; **p < 0.01, ****p < 0.0001 (one-way ANOVA).

The decomposition behavior of **HPD1** was further characterized by ^1^H NMR and HPLC. ^1^H NMR analysis confirmed the PLE-triggered decomposition of **HPD1** (**Fig. 1B**). Prior to PLE addition, **HPD1** showed characteristic signals at 2.0 ppm (proton a) and 1.5 ppm (proton b). Within 5 min of incubation at 37 °C, new signals appeared at 1.9 ppm (a’) and 2.1 ppm (b’’), matching authentic standards of acetic acid and acetaldehyde, respectively. Monitoring the decay of these signals by ^1^H NMR yielded a decomposition half-life of 15.69 ± 2.33 min for **HPD1** in the presence of 10 U PLE, with a corresponding rate constant *k* = 0.0442 ± 0.0066 min^-1^ and good linear fit (R^2^ = 0.9577), supporting pseudo-first-order kinetics (**Fig. 1C**). A transient signal at 1.2 ppm (b’) was tentatively assigned to the methyl group of the proposed hydroxymethyl persulfide intermediate, supporting the envisioned decomposition pathway (**Fig. 1B** and **1D**). HPLC analysis in phosphate buffer (pH 7.4, 37 °C) with 1 U PLE gave a decomposition half-life of 17.94 ± 2.16 min and a corresponding rate constant *k* = 0.0393 ± 0.0024 min^−1^ also following pseudo-first-order kinetics (R^2^ = 0.9893, **Fig. S1B**) Moreover, **HPD1** remained stable in PBS without PLE under the same conditions (**Fig. S1C**). Notably, **HPD1** exhibited a short half-life of within 1 min in mouse plasma (**Fig. S1D**), indicating rapid esterase-triggered activation. Such fast conversion is desirable for achieving quick onset of action, while limited systemic accumulation may enhance its safety.

### 2.3 HPD1 Alleviates Neuropathic Pain with a Superior Safety Profile

We first evaluated the analgesic efficacy of **HPD1** (14 mg/kg, i.p.) in two neuropathic pain models. In the PTX-induced neuropathy model (**Fig. 2A**, **2B** and **S2A**), **HPD1** significantly alleviated both mechanical allodynia and cold hypersensitivity. As early as 15 min post-injection, mechanical withdrawal thresholds (von Frey test) were improved, with the area under the curve (AUC) increasing from 18.70 ± 0.62 to 32.25 ± 1.15 (P < 0.0001) after **HPD1** treatment. Moreover, withdrawal response duration (acetone test) was markedly reduced, as reflected by a decrease of AUC from 65.92 ± 0.73 to 44.28 ± 0.87 (P < 0.001). These effects lasted for approximately 3 h. To rule out potential contributions from byproducts, **HPD1** was pre-incubated *in vitro* to allow complete H_2_S_2_ release, and the resulting post-release mixture was then administered to PTX-treated mice. This mixture failed to produce any analgesic activity (**Fig. S2C**, **S2D**), indicating that the antinociceptive effect of **HPD1** is specifically attributable to its H_2_S_2_-releasing capacity rather than to its byproducts. Consistently, in the CCI model (**Fig. 2C**, **2D** and **S2B**), **HPD1** produced rapid and sustained anti-allodynic effects. Mechanical allodynia was significantly attenuated, with the AUC increasing from 18.45 ± 0.69 to 33.32 ± 1.20 (P < 0.001). Attenuation of cold hypersensitivity was also observed, as indicated by a reduction of AUC from 62.55 ± 0.61 to 43.91 ± 0.69 (P < 0.0001). Collectively, these data demonstrate that **HPD1** effectively alleviates both mechanical and cold allodynia across two distinct neuropathic pain models, and that this effect is specifically mediated by its H_2_S_2_-releasing capacity.

**Figure 2.**
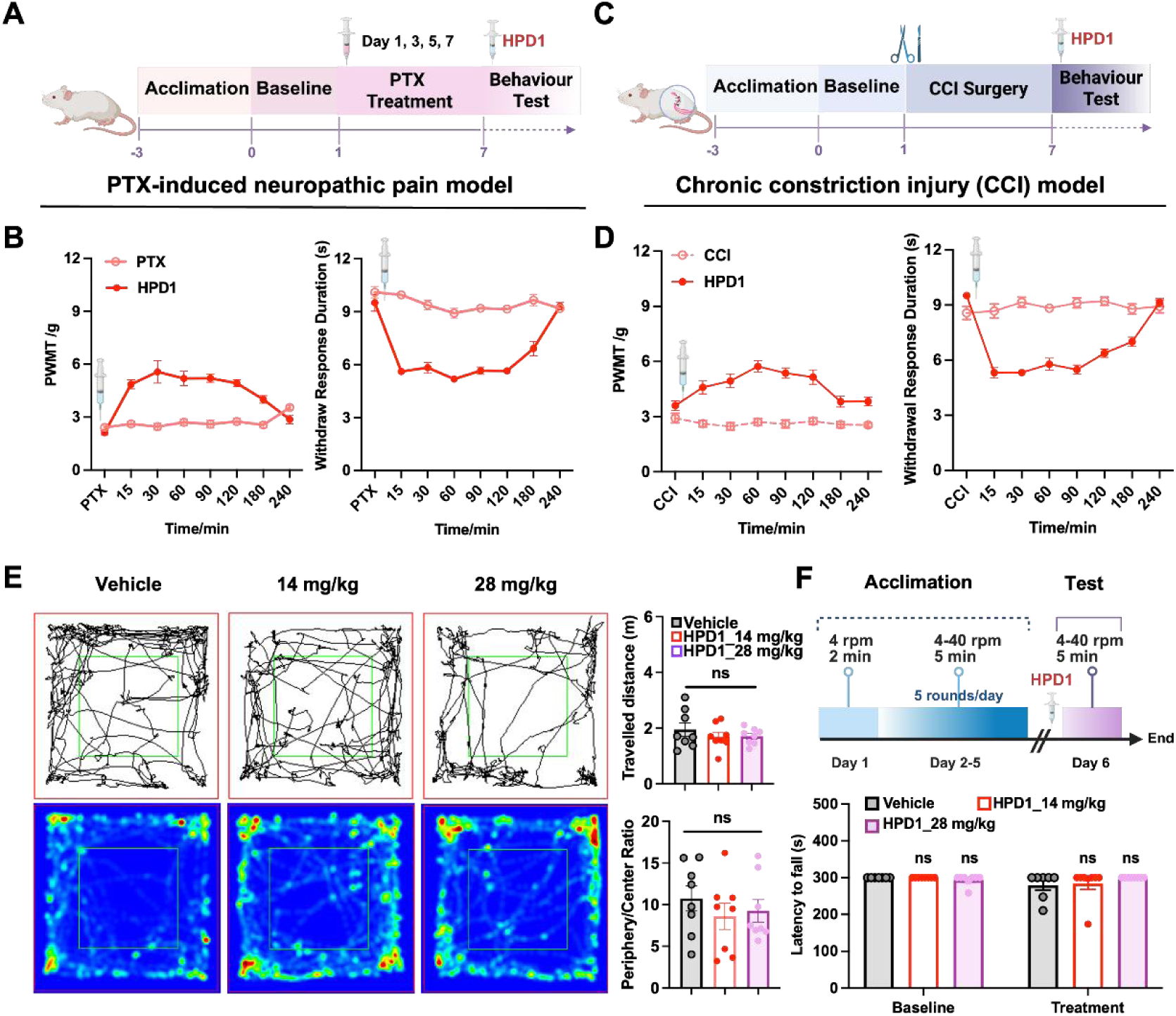
HPD1 exerts analgesic effects in chronic neuropathic pain models without affecting neurobehavioral functions. Schematic timelines of the PTX-induced neuropathy model (A) and CCI model (C). Time-course effects of **HPD1** on mechanical (left, PWMT) and cold (right, withdrawal response duration) allodynia in the PTX (B) and CCI (D) models. (E) Left panels: Representative locomotor trajectories (top) and corresponding heatmaps of locomotor activity (bottom) of mice treated with vehicle, 14 mg/kg, and 28 mg/kg **HPD1**. Right panels: Quantification of total traveled distance (top) and periphery/center ratio (bottom). (F) Setup and timeline of the rotarod test (top) and latency to fall at baseline and after **HPD1** administration (bottom). Data are mean ± SEM (n = 8 per group). One-way ANOVA followed by Tukey’s multiple comparisons test. ns, not significant. Figure 2A, 2C and 2F (Top) were created in BioRender. Kang, T. (2026) https://BioRender.com/t56dp8v.

Neurobehavioral safety assessment demonstrated that **HPD1** did not induce significant adverse effects. In the open-field test, neither total distance travelled (a measure of general locomotor activity) nor the periphery/center ratio (an indicator of anxiety-like behavior) showed significant differences among vehicle-treated control mice and mice administered therapeutic (14 mg/kg) or high (28 mg/kg) doses of **HPD1** (**Fig. 2E**). In addition, the rotarod test revealed no significant alterations in fall latency among the three groups (**Fig. 2F**). In parallel, **HPD1** demonstrated a favorable systemic safety profile. Blood biochemical analysis revealed that critical parameters, including liver function markers (ALT, AST), renal function indicators (CREA), and metabolic profiles, remained within normal ranges at 7 days, and 14 days post-administration (**Fig. 3A**). Histopathological examination of major organs (heart, liver, lung, and kidney) showed intact tissue architecture with no overt pathological abnormalities at both time points (**Fig. 3B**). Moreover, **HPD1** did not significantly affect the activities of major cytochrome P450 isoforms (**Table S2**), exhibited minimal hERG channel inhibition (IC_50_ > 30 μM, **Fig. S3**), and possessed a high median lethal dose (LD_50_ = 271.08 mg/kg in mice; **Table S3**). Collectively, these findings establish a favorable and reliable safety profile for **HPD1**.

**Figure 3.**
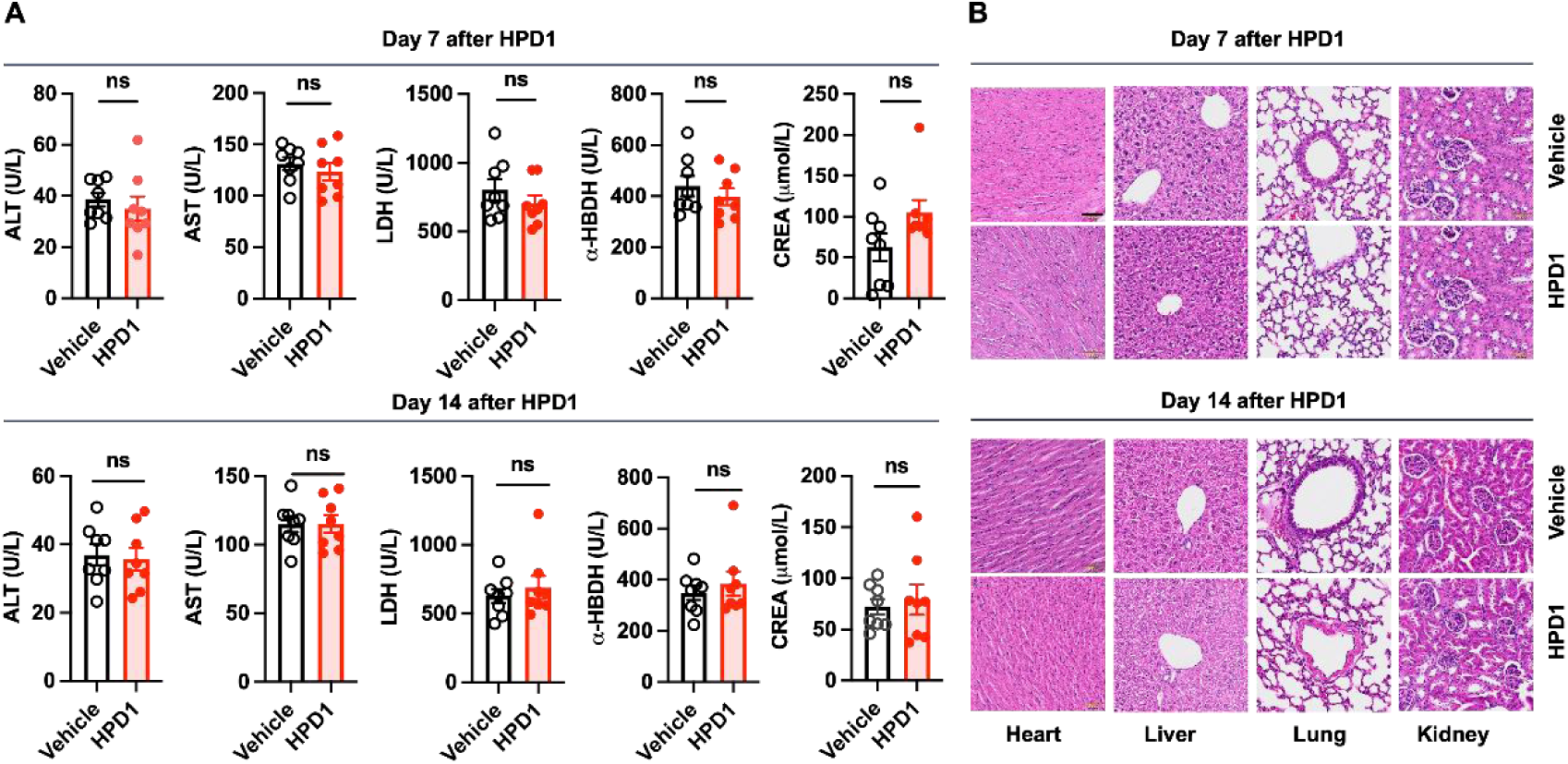
Blood biochemical and histopathological assessment of HPD1 in mice following a single high-dose administration. (A) Serum biochemical parameters, including liver function markers (ALT, AST), renal function indicators (CREA), and metabolic profiles, measured on days 7 and 14 post-treatment (n = 8). (B) Representative hematoxylin and eosin (H&E) staining of heart, liver, lung, and kidney tissues on Day 7, and Day 14 post-treatment (scale bar: 50 μm). Data are presented as mean ± SEM. Statistical analyses were performed using two-tailed unpaired Student’s t-test. ns, not significant.

### 2.4 HPD1 Normalizes PTX-Induced Hyperexcitability by Reversing A-Type Potassium Current Suppression

To investigate the effect of **HPD1** on neuronal excitability, whole-cell patch-clamp experiments on isolated DRG neurons were performed. As shown in **Fig. 4A-4C**, PTX treatment induced robust hyperexcitability compared with sham operation, as evidenced by a marked increase in action potential (AP) firing frequency (8.10 ± 1.14 Hz *vs.* 2.38 ± 0.22 Hz, P < 0.0001) and a significant reduction in rheobase current (53.23 ± 7.41 pA *vs.* 129.10 ± 17.17 pA, P < 0.05). These pathological changes were effectively reversed by **HPD1** treatment, restoring the AP firing frequency to 5.07 ± 0.83 Hz (P < 0.05) and rheobase to 125.20 ± 15.54 pA (P < 0.05) In contrast, **HPD1** did not alter AP threshold, resting membrane potential, or peak amplitude (**Fig. 4D-4F**).

**Figure 4.**
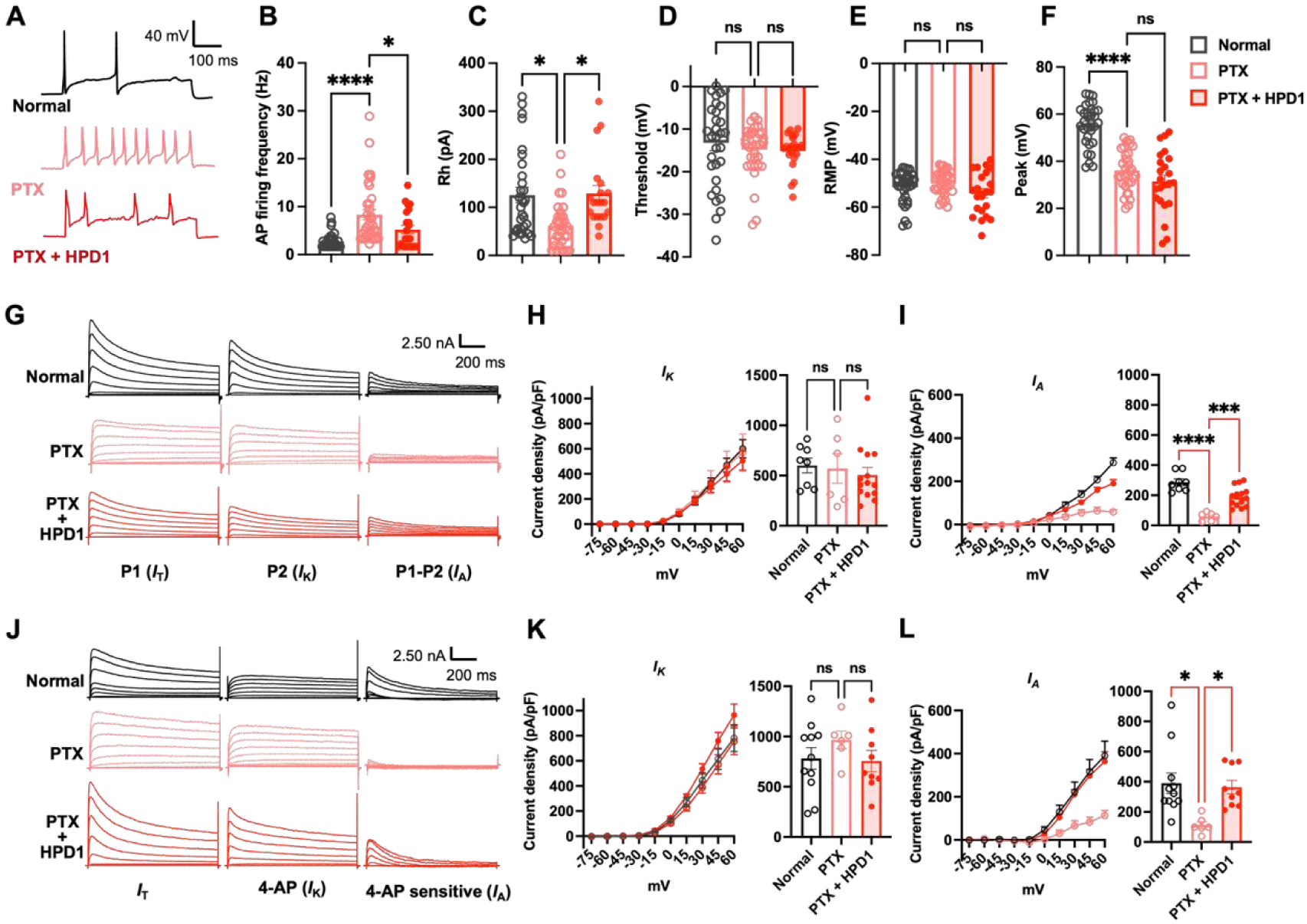
HPD1 reverses PTX-induced neuronal hyperexcitability in DRG neurons by selectively restoring the impaired *I*_A_. (A) Representative AP traces from Normal and PTX-treated DRG neurons before and after **HPD1** incubation. (B-F) Quantitative AP parameters: firing frequency (B), rheobase (C), threshold (D), rest membrane potential (RMP) (E), and peak amplitude (F) for Normal, PTX, and PTX + **HPD1** groups. (G) Two-step voltage protocol for isolating *I*_A_ (calculated as P1 (*I*_T_) *–* P2 (*I*_K_)). Representative traces from Normal, PTX, and PTX + **HPD1** neurons. (H-I) I-V relationships (left) and peak current density at +60 mV (right) for *I*_K_ (H) and *I*_A_ (I) across three groups, measured by the two-step protocol. (J) Pharmacological isolation of *I*_A_ and the representative traces using 25 mM 4-AP (*I*_T_ in the absence, *I*_K_ in the presence of 4-AP; *I*_A_ = *I*_T_ − *I*_K_). (K-L) I-V curves (left) and peak current density at +60 mV (right) for *I*_K_ (K) and *I*_A_ (L) in three groups, measured using the 4-AP inhibition method. Data are presented as mean ± SEM. Statistical analyses were performed using one-way ANOVA followed by Tukey’s multiple comparisons test. ns, not significant; *p < 0.05, ***p < 0.001, ****p < 0.0001.

We next sought to identify the ion channels mediating **HPD1**’s effect on neuronal excitability. Given the well-established role of voltage-gated sodium and calcium channels in regulating DRG neuron firing^25^, we first examined whether **HPD1** modulates these currents. However, voltage-gated sodium and calcium current densities remained unaffected following **HPD1** treatment in PTX-exposed neurons (**Fig. S4**), suggesting that these channels are not primary targets of **HPD1**. In contrast, *I*_A_ mediated by specific potassium channels is known to be suppressed in neuropathic pain models and plays a critical role in controlling firing frequency^26^. We therefore hypothesized that **HPD1** may restore *I*_A_. With a well-established two-step voltage protocol ^27,28^, total Kv currents (*I*_T_) can be evoked by protocol 1 (P1: holding potential of-120 mV, depolarization pulses from-75 mV to 60 mV in 15-mV steps). *I*_K_ can be evoked by protocol 2 (P2: holding potential of-45 mV, depolarization pulses from-45 mV to 60 mV in 15-mV steps). *I*_A_ is then obtained by subtracting P2 from P1 (**Fig. 4G**). In neurons from PTX-treated mice, the outward potassium current waveform was markedly distorted, with a clear reduction in the peak amplitude and slowed decay kinetics of the transient *I*_A_ component. Correspondingly, *I*_A_ current density was significantly decreased at +60 mV from 286.58 ± 21.94 pA/pF in normal to 58.51 ± 11.07 pA/pF in PTX. Furthermore, the proportion of *I*_A_ relative to *I*_T_ was significantly reduced in PTX-treated neurons (10.74 ± 1.89%) compared with normal controls (33.94 ± 4.00%), while the *I*_K_ proportion increased accordingly (normal: 66.06 ± 4.00%; PTX: 89.26 ± 1.89%). Notably, **HPD1** exposure effectively reversed these changes, restoring the normal waveform of *I*_A_ and recovering its current density to 190.09 ± 17.51 pA/pF, and normalizing the proportion of *I*_A_ to 30.21 ± 3.19% (**Fig. 4I**). By contrast, *I*_K_ remained stable across all experimental groups at the same potential (**Fig. 4H**). To confirm this specific effect on *I*_A_, we applied 4-aminopyridine (4-AP) to selectively block the transient potassium component in a pharmacological isolation approach (**Fig. 4J**). Quantification at +60 mV confirmed that *I*_A_ peak current density was significantly reduced in PTX-treated neurons (PTX:113.24 ± 23.11 pA/pF; normal: 390.10 ± 67.64 pA/pF), and this deficit was fully rescued by **HPD1** (363.93 ± 45.04 pA/pF), with no detectable influence on *I*_K_ (**Fig. 4K**, **4L**). In line with the voltage-protocol findings, the proportion of *I*_A_, was also decreased in PTX neurons (9.96 ± 1.16%) compared with normal controls (34.81 ± 4.17%), and HPD1 restored it to 33.97 ± 3.57%. Together, these two independent approaches consistently demonstrate that **HPD1** specifically restores the pathologically suppressed *I*_A_ in PTX-treated DRG neurons.

*I*_A_ in primary sensory neurons are predominantly mediated by Kv4 family α-subunits^29^. To further dissect the molecular basis of **HPD1**-mediated *I*_A_ potentiation, we first examined the protein expression of Kv4.1, Kv4.2, and Kv4.3 in DRG tissues from two neuropathic pain models. In mice subjected to paclitaxel treatment, **HPD1** selectively upregulated the expression of Kv4.3 without affecting Kv4.1 or Kv4.2. Meanwhile, no changes in any of the three Kv4 subtypes were observed in the CCI model (**Fig. S5**). The observation that **HPD1** upregulated Kv4.3 only in the PTX model, but not in the CCI model, indicated that the potentiation of *I*_A_ was unlikely to be driven solely by changes in Kv4 expression. Moreover, **HPD1** failed to directly potentiate Kv4.2 or Kv4.3 currents in heterologous expression systems (**Fig. S6**). Therefore, we conclude that **HPD1** acts through an indirect mechanism to augment *I*_A_, rather than as a direct agonist of Kv4 α-subunits. This indirect action may involve post-translational modifications, channel trafficking, or auxiliary subunit regulation.

### 2.5 HPD1 Exerts Analgesic Effects via the Activation of LIMK1-Cofilin Pathway

When exploring the molecular mechanism of **HPD1** using cell lines, we found that treatment with 50 μM **HPD1** for 1 h rapidly triggered morphological changes in both SH-SY5Y and ND7/23 cells. Importantly, these changes were reversible in ND7/23 cells, as the cell morphology largely recovered within 5 h, while in SH-SY5Y cells the changes lasted considerably longer (**Fig. S7**). These findings led us to hypothesize that **HPD1** may modulate actin dynamics. Phalloidin staining revealed that **HPD1** altered the normal F-actin organization, leading to a less spread and more condensed cytoskeletal pattern compared to control cells (**Fig. 5A**). Given that the actin-depolymerizing factor cofilin plays a central role in F-actin severing and turnover, and that its activity is negatively regulated by LIMK-mediated phosphorylation at Ser3^30^, we next examined whether **HPD1** modulates the LIMK-cofilin signaling axis. Western blot analysis showed that treatment with **HPD1** for 1.5 h significantly increased the phosphorylation of LIMK1 at Thr508 and its downstream substrate cofilin at Ser3 in SH-SY5Y cells, indicating pathway activation. Pretreatment with the selective LIMK1/2 inhibitor BMS-5^31,32^ suppressed **HPD1**-induced phosphorylation of both LIMK1 and cofilin (**Fig. 5B**, **5C**), confirming that **HPD1** acts upstream of this signaling cascade.

**Figure 5.**
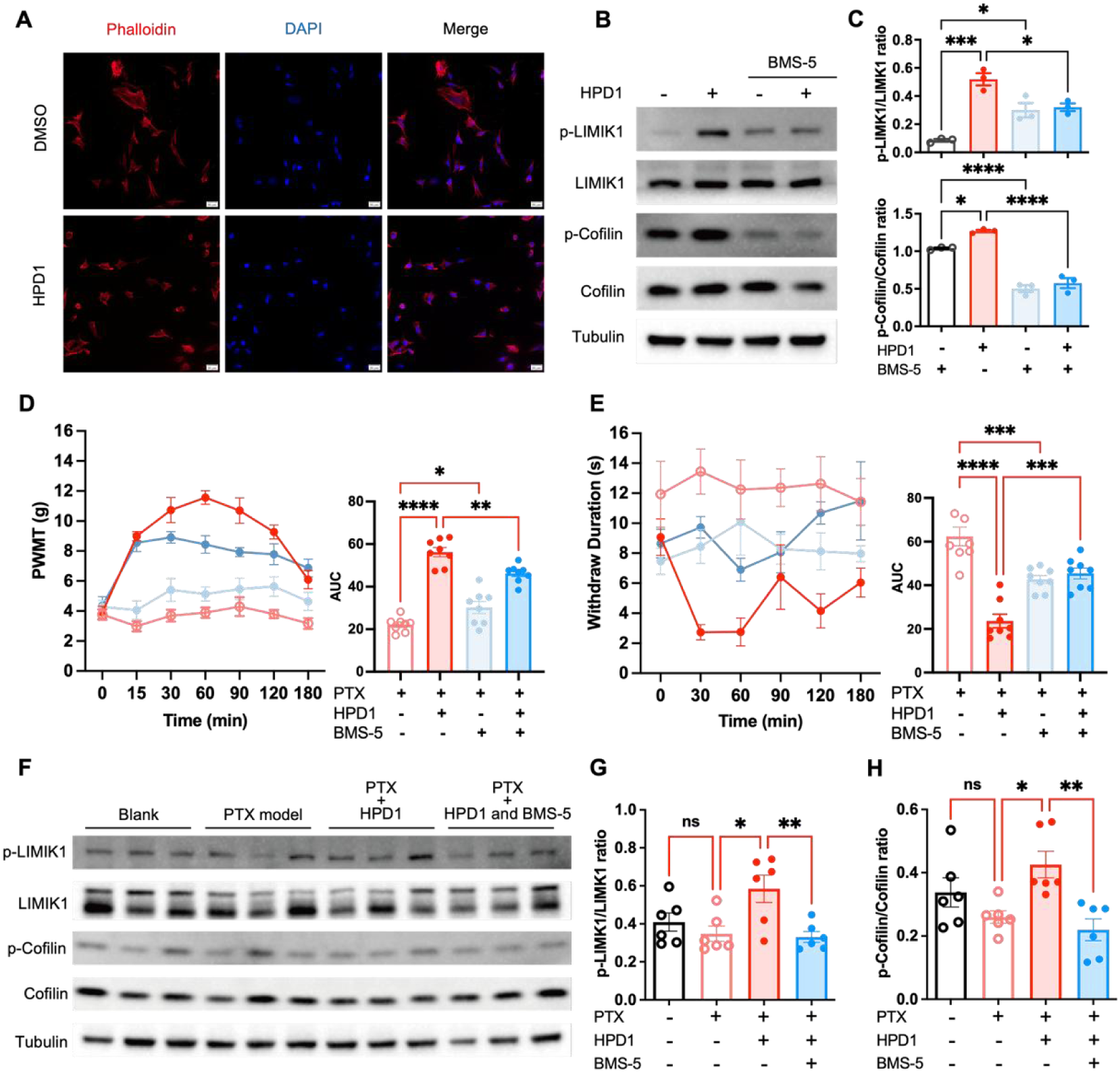
HPD1-induced analgesia is dependent on the activation of the LIMK1-Cofilin signaling pathway. (A) Phalloidin (red) and DAPI (blue) fluorescence staining of SH-SY5Y cells treated with DMSO or 50 μM **HPD1** for 1.5 h (scale bar: 20 μm). (B, C) Representative western blot images (B) and quantification (C) of p-LIMK1, LIMK1, p-Cofilin and Cofilin in SH-SY5Y cells treated with 50 μM **HPD1** for 1.5 h in the presence or absence of BMS-5 (10 μM, 4 h before **HPD1**), n = 3. (D, E) PTX-treated mice were divided into four groups: PTX alone, PTX + **HPD1**, PTX + BMS-5, and PTX + BMS-5 + **HPD1**. BMS-5 (1 mg/kg) was administered intraperitoneally 30 min prior to **HPD1** (14 mg/kg, i.p.), followed by behavioral tests. Mechanical allodynia (D) and cold allodynia (E) were assessed at indicated time points after **HPD1** administration. n = 7-8. (F-H) Representative western blot images (F) and quantification (G, H) of p-LIMK1, LIMK1, p-Cofilin and Cofilin in DRG tissues from mice subjected to the following treatments: Blank, PTX model, PTX + **HPD1**, and PTX + BMS-5 + **HPD1**. BMS-5 was administered 30 min before **HPD1**, n = 6. Data are presented as mean ± SEM. All group comparisons were performed using ordinary one-way ANOVA followed by Tukey’s multiple comparisons test. ns, not significant; *p < 0.05, **p < 0.01, ***p < 0.001, ****p < 0.0001.

We next assessed whether this pathway mediates the analgesic effects of **HPD1** using the PTX-induced neuropathic pain model. Behavioral tests revealed that **HPD1** administration significantly alleviated mechanical allodynia and cold hypersensitivity in PTX-treated mice. However, the analgesic effect of **HPD1** was markedly attenuated by pretreatment with BMS-5 (**Fig. 5D**, **5E**), indicating that LIMK1 activation is required for **HPD1** induced pain relief. To further validate the involvement of the LIMK1-cofilin pathway in the analgesic effect of **HPD1**, we performed western blot analysis of DRG tissues harvested 1 h after **HPD1** administration from PTX-treated mice. Consistent with our *in vitro* findings, **HPD1** increased the phosphorylation of LIMK1 and cofilin in DRG tissues of PTX-treated mice, and this effect was blocked by BMS-5 pretreatment (**Fig. 5F-5H**). Taken together, these data demonstrate that **HPD1** exerts analgesic effects by activating the LIMK1-cofilin pathway, which may contribute to the modulation of pain-related ion channels and neuronal excitability, potentially via actin cytoskeleton reorganization.

### 2.6 HPD1 Enhances the Filamin A-Kv4.2 Interaction and Increases A-type Potassium Currents

Having established that **HPD1** activates the LIMK1-cofilin pathway and promotes actin cytoskeleton reorganization, we next sought to identify downstream effectors that link the cytoskeleton to membrane ion channels. Filamin A, a major actin-crosslinking protein, organizes the actin network into orthogonal networks and serves as a platform for recruiting or anchoring membrane proteins^33,34^. Previous studies have reported that Filamin directly interacts with Kv4.2. This interaction targets Kv4.2 to cellular specializations, stabilizes its expression on the plasma membrane, and consequently enhances the *I*_A_ mediated by Kv4.2^16^. We therefore examined whether **HPD1** promotes the interaction between Filamin A and Kv4.2. In co-immunoprecipitation (Co-IP) experiments using the Kv4.2 antibody for pull-down, we observed a basal interaction between Kv4.2 and Filamin A, and incubation with **HPD1** for 1.5 h markedly enhanced this association (**Fig. 6A**). Western blot experiments unexpectedly revealed that BMS-5 at 10 μM for 4 h not only inhibited the LIMK-cofilin pathway but also reduced the protein abundance of Filamin A. In contrast, **HPD1** at 50 μM for 1.5 h did not significantly alter Filamin A levels, despite activating LIMK1 signaling (**Fig. 6B, 6C** and **5B**). It is plausible that Filamin A levels are sensitive to LIMK-cofilin inhibition, possibly through accelerated degradation, whereas LIMK1 activation by **HPD1** may not be sufficient to upregulate Filamin A during the short period of 1.5 h. This observation was recapitulated in DRG tissues from PTX-treated mice. Administration of **HPD1** at 14 mg/kg for 1 h had no effect on Filamin A protein expression in these tissues, whereas pretreatment with BMS-5 (30 min prior to **HPD1**) significantly reduced Filamin A abundance (**Fig. 6D**, **6E**). Thus, the reduction in Filamin A abundance by BMS-5 may represent a potential mechanism by which it could modulate the Filamin A-Kv4.2 interaction.

**Figure 6.**
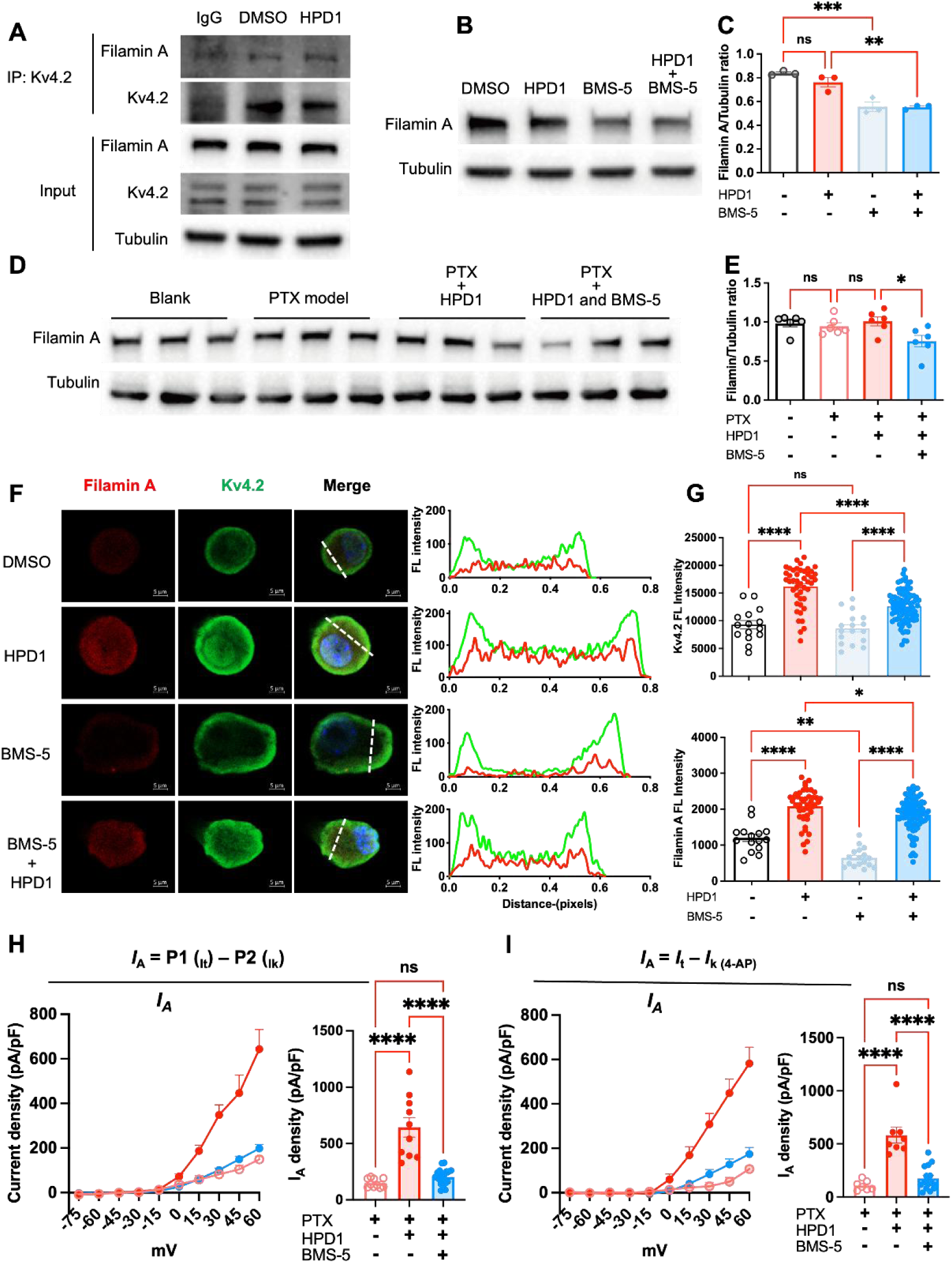
HPD1 enhances Filamin A-Kv4.2 interaction and restores A-type potassium currents via a Filamin A-dependent mechanism. (A) Lysates from SH-SY5Y cells treated with DMSO or HPD1(50 µM, 1.5 h)were immunoprecipitated with Kv4.2 antibody and immunoblotted with Filamin A and Kv4.2 antibody. (B, C) Representative western blot images (B) and quantification (C) of Filamin A in SH-SY5Y cells treated with 50 μM HPD1 for 1.5 h in the presence or absence of BMS-5 (10 μM, 4 h before HPD1), n = 3. (D, E) Representative western blot images (D) and quantification (E) of Filamin A in DRG tissues from mice subjected to the following treatments: Blank, PTX model, PTX + HPD1, and PTX + BMS-5 + HPD1. BMS-5 was administered 30 min before HPD1, n = 6. (F) Double immunostaining of Filamin A (red) and Kv4.2 (green) in primary DRG neurons from PTX-treated mice. Neurons were exposed to DMSO or 50 μM HPD1 for 1.5 h in the presence or absence of BMS-5 (10 μM, 4 h before HPD1). Profile plots were utilized to delineate the colocalization of Filamin A and Kv4.2 (scale bar: 20 μm). (G) Quantification of Filamin A and Kv4.2 fluorescence intensity in primary DRG neurons corresponding to the images in (D). (H, I) Patch-clamp recordings of A-type potassium currents (*I*_A_). The two-step voltage protocol (H) and 4-AP subtraction method (I) were applied to analyze the effects of BMS-5 pretreatment on HPD1-regulated *I*_A_ in PTX-treated neurons. Left: I-V curves; right: peak *I*_A_ density at +60 mV. Data are presented as mean ± SEM. Statistical analyses were performed using one-way ANOVA followed by Tukey’s multiple comparisons test for all other assays. ns, not significant; *p < 0.05, **p < 0.01, ***p < 0.001, ****p < 0.0001.

We next isolated primary DRG neurons from PTX-treated mice and performed double immunostaining for Filamin A and Kv4.2. Exposure to **HPD1** for 1.5 h increased the fluorescence intensity of both Filamin A and Kv4.2 in DRG neurons, and this effect was attenuated by pretreatment with BMS-5. When administered alone, BMS-5 significantly reduced Filamin A fluorescence intensity, whereas Kv4.2 intensity remained unchanged. Profile plot analysis revealed colocalization of the two proteins, which was enhanced by **HPD1**. Notably, BMS-5 diminished this colocalization, likely by reducing Filamin A abundance (**Fig. 6F**, **6G**). Finally, we performed whole-cell patch-clamp recordings to measure *I*_A_ in DRG neurons, in order to examine whether BMS-5 suppresses the **HPD1**-restored *I*_A_ density as a functional consequence of this interaction. **HPD1** application significantly increased *I*_A_ density and restored its normal current trace morphology. However, pretreatment with BMS-5 almost abolished these effects, reducing *I*_A_ density and distorting the current trace back to a pattern similar to that observed in the PTX model (**Fig. 6H**, **6I** and **S8**).

Collectively, these findings indicate that **HPD1** promotes the Filamin A-Kv4.2 interaction, likely via a LIMK/cofilin-dependent mechanism. Moreover, they support a model in which Filamin A, by enhancing Kv4.2-mediated *I*_A_, reduces DRG neuron excitability, ultimately contributing to the analgesic effect of **HPD1**.

## 3 Discussion

The management of neuropathic pain remains a major clinical challenge, highlighting the urgent need for new analgesics with distinct mechanisms of action and improved safety profiles. In this study, we introduced a novel H_2_S_2_ donor, **HPD1**, which exhibits analgesic effects in preclinical models of neuropathic pain through a previously unrecognized mechanism. **HPD1** was engineered with a rational esterase-triggered scaffold. This design ensures controlled and efficient H_2_S_2_ generation under physiological conditions. The rapid onset and sustained effects against allodynia in the CCI and PTX models further underscore its clinical potential. Mechanistically, we discovered **HPD1** activates the LIMK1-cofilin pathway, a master regulator of actin dynamics. This activation leads to cofilin inactivation and stabilization of the F-actin network, providing a more robust scaffold for Filamin A to anchor and interact with Kv4.2. Consequently, **HPD1** enhances Kv4.2 function and reverses the pathological suppression of *I*_A_, thereby alleviating pain. The critical role of this pathway was confirmed using the LIMK1/2 inhibitor BMS-5, which prevented the **HPD1**-induced increase in Filamin A-Kv4.2 co-localization, the functional restoration of *I*_A_ and the analgesic effects.

This mechanistic framework is both novel and integrative. It connects the biology of reactive sulfur species to cytoskeletal dynamics and ion channel function, offering a new paradigm for pain relief. The restoration of *I*_A_ by stabilizing the Filamin A-Kv4.2 interaction provides a powerful means to dampen the neuronal hyperexcitability that drives neuropathic pain. It is noteworthy that this mechanism is distinct from traditional analgesics that target G-protein coupled receptors or directly block voltage-gated ion channels, which may explain the favorable side effect profile of HPD1^35,36^.

Although our data implicate LIMK1 in **HPD1** induced analgesia, we did not examine LIMK2 expression or phosphorylation. LIMK2 shares high homology with LIMK1 and can also phosphorylate cofilin^37^. Given that BMS-5 inhibits both isoforms, we cannot exclude the possibility that LIMK2 contributes to the observed effects. Future studies using isoform specific genetic tools are needed to dissect the individual roles of LIMK1 and LIMK2. Furthermore, while we focused on Kv4.2 as the primary Filamin A-binding partner, the PTPP motif in the C-terminus of Kv4.2, which is required for Filamin interaction, is conserved among Kv4 family members^16^. It is therefore conceivable that Kv4.1 and Kv4.3 also participate in this interaction, and that **HPD1** may similarly enhance their association with Filamin A. Direct biochemical validation of these possibilities represents an important direction for future research.

The observation that **HPD1** activates LIMK1 to alleviate neuropathic pain seems at odds with previous evidence that pharmacological or genetic inhibition of LIMK1/2 attenuates pain in various chronic pain models^22,38^. We offer several possible explanations. First, the timing of treatment may also matter. Yang et al. reported that LIMKs are transiently activated during the development phase of chronic pain^22^. Whereas most previous studies administered LIMKs inhibitors before or simultaneously with pain induction, **HPD1** was given after neuropathy had already developed. Second, the downstream effectors engaged by LIMKs may differ across conditions. Some studies linked LIMKs to pronociceptive channels (e.g., TRPV1)^21^, whereas we identify an anti-nociceptive effector (Filamin A-Kv4.2 interaction enhancing *I*_A_). Collectively, these considerations indicate that our results do not contradict previous literature but rather expand the understanding of LIMKs signaling in pain processing.

Having established this novel mechanism, we recognize that several issues still need to be addressed. First, the precise molecular target of H_2_S_2_ released from **HPD1** is yet to be identified. H_2_S_2_ is known to mediate protein persulfidation, a post-translational modification of cysteine residues^39^. It is likely that **HPD1** promotes the persulfidation of key cysteine residues on LIMK1, cofilin, Filamin A, or an upstream regulator, thereby activating the pathway. Future studies using proteomic approaches to identify persulfidated targets following **HPD1** treatment will be crucial. Second, while our data clearly show a functional outcome in the PTX model, it will be important to test the efficacy of **HPD1** in other neuropathic pain models, such as CCI or diabetic neuropathy, to determine the generality of this mechanism. Third, we unexpectedly observed that the LIMK1/2 inhibitor BMS-5 reduced Filamin A protein levels. This effect may occur through multiple mechanisms. On one hand, inhibition of LIMK leads to cofilin-mediated actin depolymerization, which could destabilize actin-binding proteins such as Filamin A and promote their degradation. On the other hand, BMS-5 may inhibit Filamin A phosphorylation, thereby reducing its stability, as supported by previous reports that phosphorylation of Filamin A at Ser2152 enhances its resistance to calpain-mediated cleavage^40,41^. Future studies are needed to determine whether LIMK1 signaling directly regulates Filamin A phosphorylation and stability, and how these mechanisms contribute to the observed changes in Filamin A abundance.

In conclusion, we have developed a safe and effective H_2_S_2_ donor, **HPD1**, that alleviates neuropathic pain in multiple preclinical models. This work not only validates H_2_S_2_ donors as a new class of analgesics but also highlights the therapeutic potential of modulating the actin cytoskeleton to strengthen the scaffolding function of Filamin for ion channels. **HPD1**, with its novel mechanism and excellent safety profile, represents a promising lead compound for the treatment of neuropathic pain.

## 4 Experimental

### 4.1 Chemical synthesis and characterization

Details of the preparation and quality control (via NMR and HRMS) of the prodrugs are described in the Supplementary Materials.

### 4.2 Stability and Solubility assays

Stability and solubility assays were performed as described previously^42^. All experiments were conducted in duplicate for each compound.

### 4.3 Persulfide Release Kinetics

**^1^HNMR analysis of NDP-NAC decomposition kinetics. HPD1** was dissolved in 100 μL of DMSO-*d_6_* and 800 μL of D_2_O was added into the mixture, and a ^1^H NMR spectrum was recorded for the t = 0 min time point. Another sample was prepared via the following scheme: **HPD1** was dissolved in 100 μL of DMSO-*d_6_* and 230 μL of D_2_O. PLE (670 μL, 1 mg/mL stock dissolved in D_2_O) was then added, and subsequent ^1^H NMR scans were performed at different time points (5, 15, 30,60 and 120 min).

**Half-live determination of 200 μ M HPD1 by HPLC analysis.** The stability and decomposition half-life of **HPD1** were measured according to a previously described method^43^. 1 mL of 30 mM **HPD1** (2% MeOH/PBS, finial Conc. 5 mM) was added to 1 mL of PBS with 1 unit/mL esterase, containing 20 μL of 60 μM camptothecin (finial Conc. 0.1 mM) as internal standard, then the mixture was gently shaked and incubated in constant temperature water bath at 37 °C. 100 μL reaction mixture was taken out at different time points (0, 2, 5, 10, 15, 20, 30, 40, 60 and 90 min) and added into a vial containing 200 μL of ACN. The mixture was mixed well and filtered after 100 μL methanol addition, finally the filtrate was subjected to HPLC analysis.

**Quantifications of H_2_S_2_ release from HPDs by fluorescent assay.** Fluorescence-based detection of H_2_S_2_ was performed according to a previously established protocol^4^. Briefly, Na_2_S_2_ was dissolved in PBS to afford 5 mM solution. HPDs were dissolved in DMSO to afford 5 mM stock solution. 80 μ L of sulfur-contained solution was added into 4 mL of PBS buffer with or without (as control) 1 unit/mL esterase, then the mixture was incubated in 37 °C water bath for 15 min, and subsequently at RT. for 5 min. Additionally 20 μL of 5 mM CTAB and 160 μL of 0.5 mM DSP-3 was added, and the reaction mixture was incubated at RT. for another 5 min. Finally, the fluorescent intensity at 515 nm of each group was recorded with excitation at 490 nm using fluorescence spectra instrument (HORIBA, iHR320).

### 4.4 Animals and Ethics Statement

All mice were housed in a pathogen-free facility under controlled temperature (23 ± 1 °C) and a 12-h light/dark cycle (lights on 07:00-19:00), with standard rodent chow and water provided ad libitum. Male ICR mice (6-8 weeks old; Chengdu Dossy Experimental Animals Co., Ltd.) were used for dorsal root ganglion (DRG) neuron isolation, culture, and electrophysiological recordings. Adult ICR mice (6-8 weeks old, with equal numbers of males and females; Chengdu Dossy Experimental Animals Co., Ltd.) were used for all other animal experiments. Mice were acclimated to the housing environment for at least 3 days. For behavioral tests requiring adaptation to the testing apparatus (e.g., rotarod), additional acclimation was performed as described in the respective sections. All animal procedures were approved by the Animal Research Committee of West China Hospital of Sichuan University (Protocol No. 20220311021) and were conducted in accordance with the NIH Guide for the Care and Use of Laboratory Animals and the ethical guidelines of the International Association for the Study of Pain. Mice were euthanized with a lethal dose of pentobarbital for tissue collection, and all efforts were made to minimize animal suffering.

### 4.5 Adminidtration

For intraperitoneal administration, **HPD1** was dissolved in a vehicle consisting of 10% DMSO, 10% PEG-400, and 80% saline, and was injected at doses of 14 or 28 mg/kg in a volume of 0.2 mL. BMS-5 was dissolved in saline containing 0.1% DMSO and administered at 1 mg/kg in the same injection volume.

### 4.6 Primary culture of dorsal root ganglion neurons

Primary cultures of dorsal root ganglion (DRG) neurons were prepared following a previously reported protocol^44–46^. Male ICR mice were anesthetized with ether and then rapidly decapitated. The DRGs from spinal L4–6 segments were removed on ice and immediately collected into a 35-mm culture dish (#706001; Nest), then digested in 2 mg/ml collagenase IV (#C4-BIOC; Sigma Aldrich) for 30 min at 37 °C, followed by 1% trypsin (#P7545; Sigma Aldrich) for another 10 min. Digested ganglia were gently washed with neurobasal medium and mechanically dissociated by passage through the pipette. The isolated DRG neurons were seeded on poly-D-lysine-coated 9 mm coverslips and cultured overnight at 37 °C in 5% CO_2_ in neurobasal medium supplemented with 2% B27 (#17504044; Gibco) and penicillin and streptomycin (100 U/mL and 100 μg/mL) for 4-8 h.

### 4.7 Electrophysiological Recordings

Whole-cell patch-clamp recordings were performed at 22.0 ± 2.0 °C. Borosilicate glass microelectrodes (2-6 MΩ) were pulled using a P1000 puller (Sutter Instruments). Recordings were conducted with a Sutter 200B amplifier in current-and voltage-clamp modes. Series resistance was compensated by ∼80%. Signals were sampled at 10 kHz, low-pass filtered at 2 kHz, and analyzed with Igor 9 (WaveMetrics) and GraphPad Prism 10.3.0. Drugs were pre-incubated for 4-8 h and continuously perfused at 10 μM via a PV-pump (ALA Scientific). Recordings were obtained from small-diameter (< 30 μm) DRG neurons within 8 h of plating, with resting membrane potentials more negative than-45 mV.

For neuronal excitability recordings, the bath solution contained (mM): 140 NaCl, 5 KCl, 2 CaCl_2_, 1 MgCl_2_ 10 HEPES, 10 D-glucose (pH 7.4, 310-315 mOsm). The pipette solution contained (mM): 126 KCl, 10 NaCl, 1 MgCl_2_, 10 EGTA, 2 Na-ATP, 0.1 Mg-GTP (pH 7.3, 300 ± 10 mOsm). Action potentials were evoked by 600-ms depolarizing current injections (0–400 pA, 10-pA increments).

For Na^+^ current (*I*_Na_) recording: the internal pipette solution contained (in mM): 120 CsCl, 10 NaCl, 10 TEA-Cl, 10 HEPES, 10 EGTA, 1 CaCl_2_, and 1 MgCl_2_ (pH 7.2, 290-310 mOsm). The external solution contained (in mM): 120 NaCl, 1 CaCl_2_, 20 TEA-Cl, 5 KCl, 1.25 MgCl_2_, 10 HEPES, and 10 D-glucose (pH 7.4, 310-315 mOsm).

Currents were elicited by a 10-ms pre-pulse at-70 mV, followed by a 20-ms depolarizing step to 0 mV, with 10-s inter-step intervals^47,48^.

Total calcium currents were measured after blocking Na^+^ and K^+^ currents with 1 μM TTX and 30 mM TEA-Cl^47,49^. The extracellular solution contained (in mM): 110 N-methyl-D-glucamine, 10 BaCl_2_, 10 HEPES, and 10 D-glucose (pH 7.3, 310-315 mOsm). The intracellular solution contained (in mM): 150 CsCl, 10 HEPES, 5 Mg-ATP, and 5 BAPTA (pH 7.3, 290-310 mOsm). Currents were evoked by a 500-ms depolarizing step to 0 mV from a holding potential of-80 mV. After perfusion with **HPD1** for at least 5 min, currents were recorded at 10-s intervals after stabilization, and average values were calculated.

Total and subtypes of potassium currents (*I*_T_, *I*_A_, *I*_K_) were measured using both pharmacological block and voltage-protocol subtraction^27,28^. The pipette solution contained (in mM): 115 potassium gluconate, 25 KCl, 1 CaCl_2_, 2 MgCl_2_, 10 EGTA, 10 HEPES, and 5 Na_2_ATP (pH 7.3, 290-310 mOsm). The bath solution contained (in mM): 140 choline chloride, 5 KCl, 2 CaCl_2_, 1 MgCl_2_, 10 HEPES, 1 CdCl_2_, and 10 D-glucose (pH 7.4, 310-315 mOsm). For pharmacological block, 25 mM TEA (*I*_K_ blocker) or 5 mM 4-AP (*I*_A_ blocker) was added to the bath, with choline chloride adjusted to 115 or 135 mM, respectively. For protocol subtraction, total K^+^ current (*I*_T_) was elicited by Protocol 1 (P1): holding potential-120 mV, steps from-75 mV to +60 mV (15-mV increments); the sustained delayed-rectifier current (*I*_K_) was elicited by Protocol 2 (P2): holding potential-45 mV, steps from-45 mV to +60 mV (15-mV increments). The A-type K^+^ current (*I*_A_) was calculated as *I*_A_ = P1 (*I*_T_) - P2 (*I*_K_).

### 4.8 Paclitaxel induced CIPN model

To establish a CIPN model, the paclitaxel injection regimen, PTX-H20053001, of the Yangtze River Pharmaceutical Group (4 ×, 2 mg/kg, i.p) was applied on day 1, 3, 5, and 7 as previously described^50^. Mice were i.p. injected with vehicle or **HPD1** on day 8, then mechanical withdrawal threshold and cold allodynia were assessed. Behavioral responses were recorded for three to four hours after administration.

### 4.9 Chronic constriction injury (CCI) mice model

Chronic constriction injury (CCI) of the sciatic nerve was induced as previously described^51,52^. ICR mice were anesthetized with isoflurane, and under aseptic surgical conditions, three ligatures of 5-0 chromic gut were loosely tied around the sciatic nerve 1 mm apart and proximal to the trifurcation. For fluid replacement, mice received 1 mL of warmed lactated Ringer’s solution subcutaneously at the end of surgery. One week after surgery, mice were dosed with **HPD1** or vehicle and tested for mechanical and cold analgesic effects post-treatment.

### 4.10 Behavioral test

For Von Frey test, mice were acclimated in Plexiglas cages (20 cm × 20 cm × 15 cm), and paw withdrawal mechanical threshold (PWMT) was measured with an electronic Von Frey apparatus (No. 2391, IITC Life Science Inc, Woodland Hills, CA) by applying vertical force to each hind paw plantar surface; mechanical sensitivity was evaluated at 0.25, 0.5, 1, 1.5, 2, 3, and 4 h post-treatment. For cold allodynia, mice were acclimated for 15 min in a transparent Plexiglas chamber on an elevated wire mesh grid. A 10-12 μL aliquot of acetone (≥ 99.5%) was gently applied to the hind paw plantar surface with a micropipette (no mechanical contact); each paw was tested twice per time point (5-min interval) with a 60-s cutoff. Nocifensive behavior (paw withdrawal, licking, shaking) duration within 1 min post-acetone application was recorded by a blinded investigator with a digital stopwatch, and the mean of two replicates was used as the cold pain sensitivity index.

### 4.11 Open field test

Locomotor activity and anxiety-like behavior were assessed using the open field test according to established protocols^53,54^. Mice were acclimated to the testing room and apparatus for 1 h daily for 3 consecutive days before the experiment. Animals were randomly divided into three groups (n = 8): vehicle control, **HPD1** 14 mg/kg, and **HPD1** 28 mg/kg. Behavioral testing was performed approximately 1 h after administration (i.p.). Each mouse was placed in the open-field chamber and recorded for 10 min using a Smart v2.5 video-tracking system (Panlab, Barcelona, Spain). The time ratio in the central zone/the peripheral zone was analyzed during the first 5 min for novelty exploration, and the total distance travelled was measured during the last 5 min for locomotor activity.

### 4.12 Rotarod test

The rotarod test was performed as previously described^55^. The rotarod test is divided into training stages and testing stages. Animals were randomly assigned to three groups (n = 8): vehicle control, **HPD1** 14 mg/kg, and **HPD1** 28 mg/kg. On the first training day, mice were trained on the rotarod (ENV-575M, MED Associates, Fairfax, VT, USA) at a constant speed of 4 rpm for 2 min. From the second training day onward, the rotarod was set to accelerate from 4 rpm to 40 rpm over 300 s, with 5 training trials per day and at least 30 min inter-trial intervals for 4 consecutive days. Mice unable to remain on the rotarod for 300 s by the end of training were excluded from subsequent test. On the test day, the accelerating protocol (4-40 rpm, 300 s) was used. Latency to fall and the corresponding rotational speed were recorded before and 1 h after drug administration.

### 4.13 Biochemical Analysis and Histopathological Examination

Mice (6-8 weeks) single administrated with vehicle or **HPD1** (28 mg/kg) were anesthetized for serum and viscus collection for blood biochemical analysis and hematoxylin staining on day 1, day 7 and day 14.

### 4.14 Western blots

For *in vitro* studies, SH-SY5Y cells were seeded in 6-well plates and treated as indicated. After treatment, cells were washed twice with ice-cold phosphate-buffered saline (PBS) and lysed in RIPA buffer (Beyotime, China) supplemented with a protease inhibitor cocktail (HY-K0010, MCE, USA). For tissue analysis, The L4-6 DRGs were rapidly harvested from anesthetized ICR mice at designated time points following drug administration. Tissues were homogenized in ice-cold RIPA buffer containing protease inhibitors using a tissue grinder. The lysates were centrifuged at 12,000 × g for 15 min at 4 °C, and the supernatants were collected. Protein concentrations were determined using a BCA Protein Assay Kit (Beyotime, China).

Equal amounts of protein were denatured in 5× loading buffer at 95 °C for 10 min, separated by 4-20% SDS-polyacrylamide gel electrophoresis (SDS-PAGE), and subsequently transferred onto polyvinylidene fluoride (PVDF) membranes (Millipore, USA) using a wet transfer system (Bio-Rad, USA) at 250 mA for 70-180 min at 4 °C.

Membranes were blocked with 5% non-fat milk or bovine serum albumin (BSA) in Tris-buffered saline containing 0.1% Tween-20 (TBST) for 1 h at room temperature. They were then incubated overnight at 4 °C with primary antibodies diluted in blocking buffer. The following primary antibodies were used: anti-Kv4.1 (A11599-1, Boster, 1:1000), anti-Kv4.2 (21298-1-AP, Proteintech, 1:1000), anti-Kv4.3 (A6927, ABclonal Technology, 1:1000), anti-p-LIMK1 (Thr508) (AF3345, Affinity, 1:1000), anti-LIMK1 (67974-1-Ig, Proteintech, 1:2000), anti-p-cofilin (Ser3) (#3313, Cell Signaling Technology, 1:1000), anti-cofilin (10960-1-AP, Proteintech, 1:2000), anti-Filamin A (67133-1-Ig, Proteintech, 1:2000), and anti-GAPDH (60004-1-Ig, Proteintech, 1:2000) / anti-α-Tubulin(66031-1-lgG,Proteintech, 1:2000) as a loading control.

After washing three times with TBST, membranes were incubated with horseradish peroxidase (HRP)-conjugated secondary antibodies for 1 h at room temperature. Following three additional washes, protein bands were visualized using an enhanced chemiluminescence (ECL) substrate (4A biotech, China) and detected with a chemiluminescence imaging system (Bio-Rad, USA).

Densitometric analysis of the bands was performed using ImageJ software (NIH, USA). The target protein expression levels were normalized to GAPDH or Tubulin, and phosphorylation levels were normalized to the corresponding total protein. All experiments were repeated at least three times independently, and representative blots are shown.

### 4.15 Co-Immunoprecipitation (Co-IP)

For *in vitro* Co-IP, SH-SY5Y cells were cultured in 10 cm dishes and treated as indicated. After treatment, cells were washed twice with ice-cold PBS and lysed in 500 μL of NP40 lysis buffer (Beyotime, China) containing protease inhibitor cocktails. All lysates were incubated on ice for 30 min, followed by centrifugation at 12,000 × g for 15 min at 4 °C. The supernatants were collected, and protein concentrations were determined using a BCA Protein Assay Kit. A small aliquot (40 μL) of each lysate was saved as “input” for subsequent Western blot analysis.

For each sample, 500 μg of total protein was incubated with 1 μg anti-Kv4.2 antibody or control IgG overnight at 4 °C with continuous rotation. The next day, 40 μL of Protein A/G Magnetic Beads (HY-K0202, MCE, USA) were added to each sample and incubated at 4 °C for an additional 4 h. After washing five times with 500 μL PBST buffer, the beads were resuspended in 40 μL of 1× SDS loading buffer and boiled at 95°C for 10 min to elute the immunoprecipitated proteins.

### 4.16 Immunofluorescence Staining and Imaging

SH-SY5Y cells were seeded on glass coverslips coated with poly-D-lysine in 24-well plates. After drug treatments, cells were washed briefly with PBS and fixed with 4% paraformaldehyde (PFA) in PBS for 15 min at room temperature. Cells were then permeabilized with 0.2% Triton X-100 in PBS for 10 min and blocked with 5% bovine serum albumin (BSA) in PBS for 1 h at room temperature. For F-actin staining, cells were incubated with YF 594-conjugated Phalloidin (YP0052, UElandy, 1:200) for 30 min at room temperature in the dark. Nuclei were counterstained with DAPI.

DRG primary cells from PTX-treated mice were seeded on glass coverslips coated with poly-D-lysine in 24-well plates. After drug treatments, cells were washed briefly with PBS and fixed with 4% paraformaldehyde (PFA) in PBS for 15 min at room temperature. Cells were then permeabilized with 0.2% Triton X-100 in PBS for 10 min and blocked with 10% goat serum in PBS for 1 h at room temperature. Then, cells were incubated overnight at 4 °C with primary antibodies: anti-Filamin A (#44873, Cell Signaling Technology, 1:200) and anti-Kv4.2 (sc-390571, Santa Cruz, 1:100). After washing three times with PBS, cells were incubated with CoraLite594 conjugated goat anti-rabbit secondary antibody (SA00013-4, Proteintech, 1:400) and BF488 conjugated goat anti-mouse secondary antibody (bs-0296G-BF488, Bioss, 1:400) for 1 h at room temperature in the dark. Nuclei were counterstained with DAPI.

Fluorescent images were captured using a confocal laser scanning microscope (LSM 880, Zeiss, Germany) with a 40× objective. For co-localization analysis, images were processed using ImageJ software (NIH, USA). The degree of co-localization between Filamin A and Kv4.2 was quantified using Pearson’s correlation coefficient with the JACoP plugin. For quantification of fluorescence intensity, regions of interest (ROIs) were drawn around individual neurons, and the mean fluorescence intensity was measured. At least 30 neurons per group from three independent experiments or animals were analyzed. All imaging parameters were kept constant across all experimental groups.

### 4.17 Data Analysis

Data were expressed as mean ± SEM. The sample size is indicated in figure legends. Statistical analyses were done with GraphPad Prism 10.3.0. (GraphPad Software, Inc., La Jolla, CA, USA), Igor 9 Software (WaveMetrics, Portland, OR, USA), and Microsoft Excel. Behavioral tests, H&E staining, blood biochemistry, and electrophysiology data were analyzed by one-way ANOVA or unpaired two-tailed Student’s t-tests. p< 0.05 was considered statistically significant.

## Supporting Information

### Materials and Methods

#### Chemistry

All solvents were of reagent grade and were purchased from Fisher Chemicals. Reagents were purchased from Aldrich, Oakwood, or VWR. ^1^H-(400 MHz) and ^13^C-(100 MHz) NMR spectra were recorded on a Bruker Avance 400 MHz NMR spectrometer. Mass spectral analyses were performed on an ABI API 3200 (ESI-Triple Quadruple) or a Waters Q-TOF Micros. High Performance Liquid Chromatography (HPLC) studies were performed on a DEGASSING UNIT DGU-20A3R.(C) system with an Ultimate® XB-C18 reverse phase column (5 μm, 4.6 mm × 250 mm) eluted at a rate of 1 mL/min with solvent A (0.1% trifluoroacetic acid in water) and solvent B (acetonitrile) for HPLC analysis, with the ratio initiating from 1:9 to 9:1 after during 12 min, and monitored at 210 nm.

### Synthesis of 1-(acetylthio)ethyl acetate (b1)

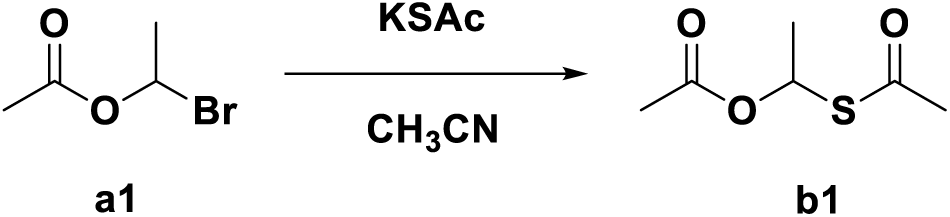

To a round bottom flask equipped with nitrogen protection was added a solution of 1-bromoethyl acetate **a1** (1.67 g, 10 mmol, 1eq) in acetone (15 mL) at 0 °C, which was added potassium ethanethioate (1.14 g, 10 mmol, 1eq), and the mixture was then stirred at room temperature overnight. The reaction mixture was filtered and the filtrate was rotary evaporated. The residue was dissolved in CH_2_Cl_2_ and washed with H_2_O. The aqueous fraction was extracted with CH_2_Cl_2_. The combined organic layers were dried over Na_2_SO_4_, filtered, rotary evaporated and then dried in vacuo. The crude extract was purified by column chromatography and the product was eluted with 0-3 % ethyl acetate in petroleum ether to give 0.5 g of the pure desired product as brown oil (yield: 31.25%). 1H NMR (400 MHz, Chloroform-d) δ 6.41 (q, J = 6.4 Hz, 1H), 2.26 (s, 3H), 1.98 (s, 4H), 1.54 (d, J = 6.0 Hz, 3H).

### Synthesis of disulfanediylbis(ethane-1,1-diyl) diacetate (HPD1)

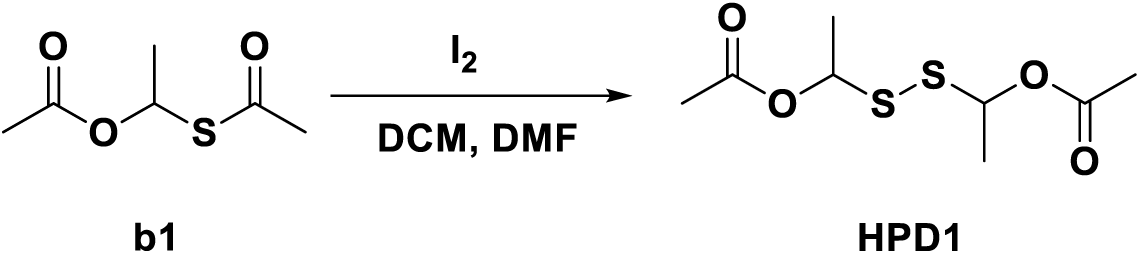

To the solution of 1-(acetylthio)ethyl acetate **b1** (2 g, 12.33 mmol, 1 eq) in MeCN (30 mL) was added I_2_ (3.13 g, 24.66 mmol, 2 eq) and NIS (1.39 g, 6.16 mmol, 0.5eq) at-10 °C. The mixture was stirred at-10 °C for 3h. The mixture was diluted with Na_2_S_2_O_3_ solution (5% in H_2_O, 100 mL) and extracted with DCM (50 mL×3). The combined organic layers were dried over Na_2_SO_4_, filtered and the filtrate was concentrated in vacuo. The residue was TLC (PE:EA = 10:3) to give the crude product. The crude was purified by Pre-HPLC to give the product **HPD1** (50 mg) as light yellow oil (yield: 3.4%). ^1^H NMR (400 MHz, Chloroform-d) *δ* ^1^H NMR (400 MHz, Chloroform-d) δ 5.96 (dd, *J* = 6.4, 4.0 Hz, 2H), 2.09 (d, *J* = 4.0 Hz, 6H), 1.58 (dd, *J* = 6.4, 6H). ^13^C NMR (101 MHz, Chloroform-d) *δ* 169.81, 78.26, 76.76, 21.07, 21.05, 20.26, 20.07. HRMS (ESI-QQQ) *m/z* [M + Na]^+^ Calculated for C_8_H_14_O_4_NaS_2_: 261.0226, found: 261.0222.

### Synthesis of 1-bromoethyl benzoate (a2)

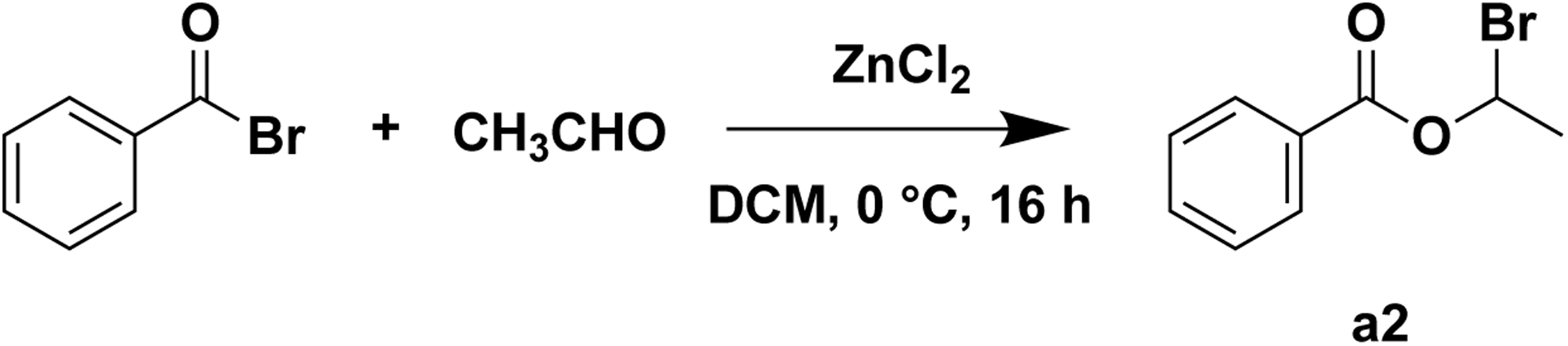

To a solution of commercially available benzoyl bromide (2.0 g, 1.0 eq.) in DCM (20 mL) was added ZnCl_2_ (74 mg, 0.05 eq.) at 0 ℃. The mixture was then stirred under N_2_ for 30 minutes. After that, acetaldehyde (381 mg, 0.8 eq.) in DCM (10 mL) was added into the mixture and stirred for 16 hours at 0 ℃. The reaction was then monitored by TLC. After completion, the mixture was quenched with NaHCO_3_ aq., then the organic layer was separated and washed with NaCl aq. for twice. After that, the organic layer was dried over Na_2_SO_4_ and concentrated in vacuo to afford a to remove the solvent away. The residue was purified by flash column chromatography (petroleum ether: ethyl acetate = 100: 1) to afford desired product as a colorless oil (1.03 g, yield: 41.8%). ^1^H NMR (400 MHz, Chloroform-*d*) δ 8.11 – 8.05 (m, 2H), 7.66 – 7.58 (m, 1H), 7.47 (t, *J* = 7.8 Hz, 2H), 6.97 (q, *J* = 5.9 Hz, 1H), 2.14 (d, *J* = 5.9 Hz, 3H).

### Synthesis of 1-(acetylthio)ethyl benzoate (b2)

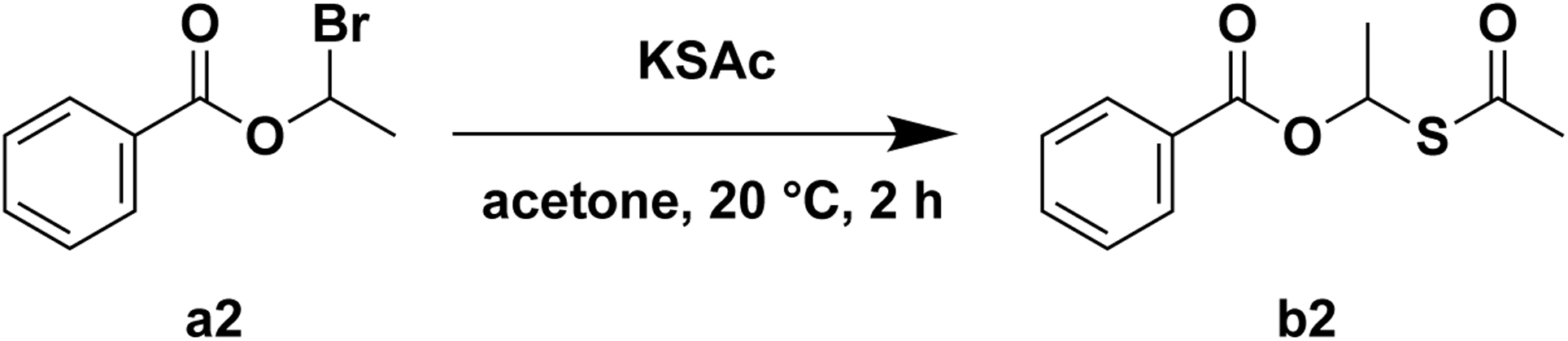

To a solution of 1-bromoethyl benzoate (1.00 g, 1.0 eq.) in acetone (20 mL) was added KSAc (498 mg, 1.0 eq.) slowly at 20 ℃. The mixture was then stirred at 20 ℃ for 2 hours. The reaction was then monitored by TLC. After completion, the mixture was filtered and the filtrate was concentrated to afford a residue. The residue was purified by flash column chromatography (petroleum ether: ethyl acetate = 10:1) to afford desired product as a colorless oil (700 mg, yield: 69.2%). ^1^H NMR (400 MHz, Chloroform-*d*) δ 7.98 – 7.92 (m, 2H), 7.52 – 7.46 (m, 1H), 7.36 (t, *J* = 7.7 Hz, 2H), 6.67 (q, *J* = 6.6 Hz, 1H), 2.27 (s, 3H), 1.69 (d, *J* = 6.7 Hz, 3H).

### Synthesis of Disulfanediylbis(ethane-1,1-diyl) dibenzoate (HPD2)

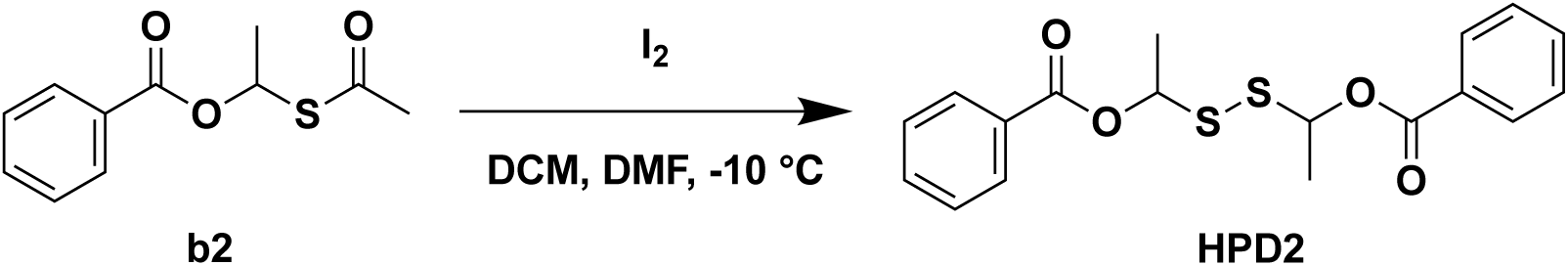

To a solution of 1-(acetylthio)ethyl benzoate (700 mg, 1.0 eq.) in DCM (10 mL) was added I_2_ (1.58 g, 2.0 eq.) in DMF (5 mL) dropwise at-10 ℃. The mixture was then stirred at-10 ℃ for 1 hour and then warmed up to room temperature and stirred for 4 hours. The reaction was then monitored by TLC. After completion, the mixture was quenched with Na_2_S_2_O_3_ aq. and extracted with ethyl acetate for three times. The organic layer was separated and dried over Na_2_SO_4_ and concentrated in vacuo to afford a residue. The residue was purified by flash column chromatography (petroleum ether: ethyl acetate = 10: 1) to afford desired product as a colorless oil (220 mg, yield: 19.4%). ^1^H NMR (400 MHz, Chloroform-*d*) δ 8.07 – 8.01 (m, 4H), 7.61 – 7.54 (m, 2H), 7.48 – 7.39 (m, 4H), 6.28 – 6.20 (m, 2H), 1.73 – 1.66 (m, 6H). ^13^C NMR (101 MHz, Chloroform-*d*) δ 165.45, 165.43, 133.37, 133.32, 129.86, 129.82, 129.63, 129.57, 128.49, 128.43, 79.10, 78.30, 20.47, 20.38 (two signals observed for most carbons due to the presence of enantiomers). HRMS (ESI-QQQ) *m/z* [M + Na]^+^ Calculated for C_18_H_18_O_4_NaS_2_: 385.0539, found: 385.0525.

### Synthesis of vinyl 2-acetoxybenzoate (a3-i)

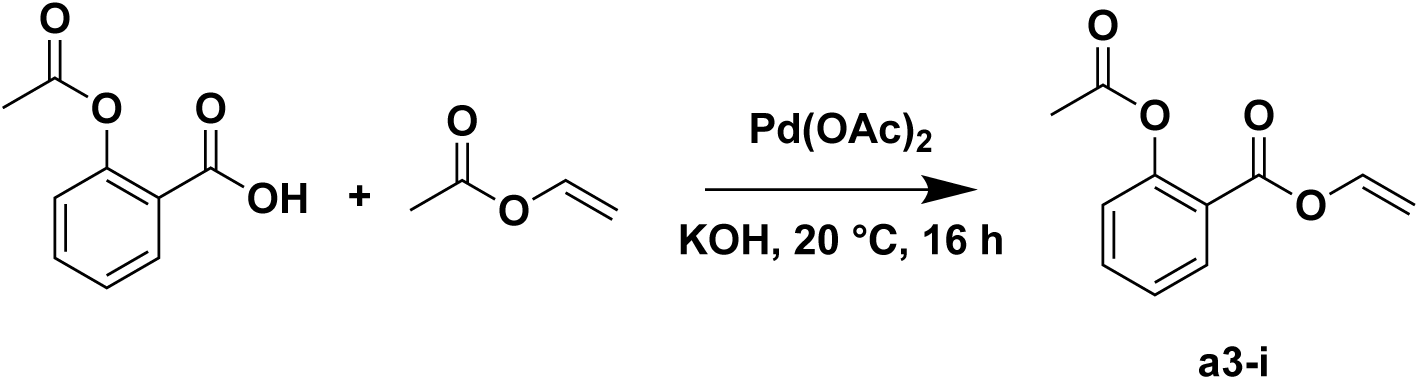

A minture of 2-acetoxybenzoic acid (500 mg, 1.0 eq.), Pd(OAc)_2_ (31 mg, 0.05 eq.) and KOH (8 mg, 0.05 eq.) in vinyl acetate (4 mL) was stirred at 20 ℃ for 16 hours. The reaction was then monitored by TLC. After completion, the reaction mixture was filtered. The filtrate was concentrated in vacuo to afford a residue. The residue was then purified by flash column chromatography (petroleum ether: ethyl acetate = 4: 1) to afford desired product as a colorless oil (300 mg, yield: 52.4%). ^1^H NMR (400 MHz, DMSO-*d*_6_) δ 8.07 – 8.02 (m, 1H), 7.79 – 7.72 (m, 1H), 7.49 – 7.44 (m, 1H), 7.43 – 7.36 (m, 1H), 7.33 – 7.27 (m, 1H), 5.13 – 5.05 (m, 1H), 4.87 – 4.77 (m, 1H), 2.30 (s, 3H).

### Synthesis of 1-bromoethyl 2-acetoxybenzoate (a3)

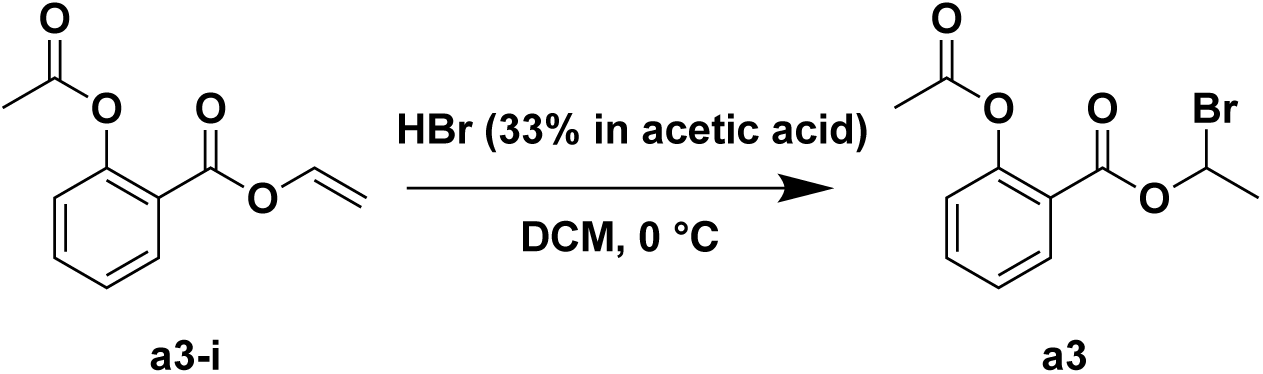

To a solution of HBr (428 mg, 33% in acetic acid, 1.2 eq.) in DCM (5 mL) was added vinyl 2-acetoxybenzoate (300 mg, 1.0 eq.) dissolved in DCM (5 mL) dropwise at 0 ℃. The mixture was then stirred at 0 ℃ for 1 hour. After completion, the mixture was poured into ice water and extracted with DCM. The organic layer was separated, dried and concentrated in vacuo to afford the crude product as a yellow oil (448 mg). The crude product was used in the next step without any purification.

### Synthesis of 1-(acetylthio)ethyl 2-acetoxybenzoate (b3)

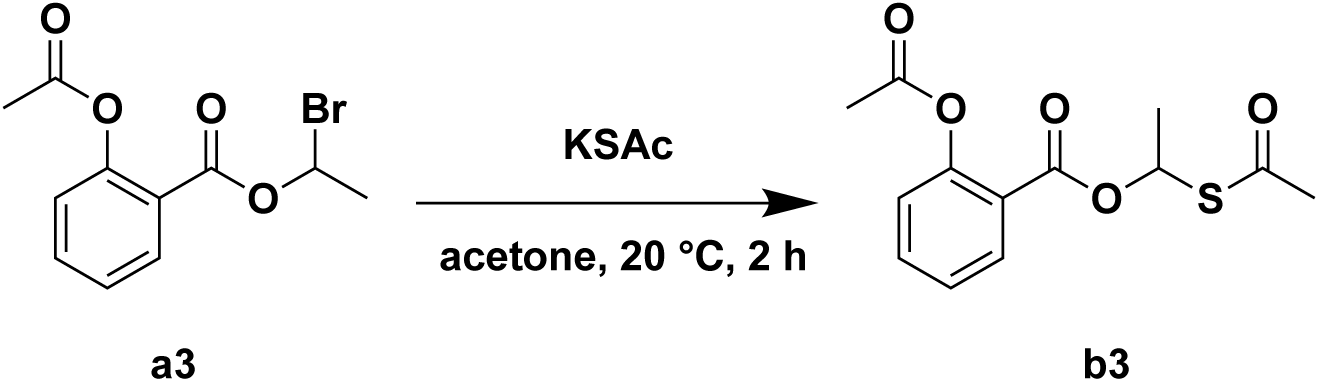

To a solution of 1-bromoethyl 2-acetoxybenzoate (448 mg, 1.0 eq.) in acetone (10 mL) was added KSAc (200 mg, 1.2 eq.) slowly at 20 ℃. The mixture was then stirred at 20 ℃ for 16 hours. The reaction was then monitored by TLC. After completion, the mixture was filtered and the filtrate was concentrated to afford a residue. The residue was purified by flash column chromatography (petroleum ether: ethyl acetate = 93: 7) to afford desired product as a colorless oil (153 mg, yield: 34.8%). ^1^H NMR (400 MHz, DMSO-*d*_6_) δ 7.93 – 7.87 (m, 1H), 7.74 – 7.66 (m, 1H), 7.45 – 7.39 (m, 1H), 7.28 – 7.22 (m, 1H), 6.56 (q, *J* = 6.6 Hz, 1H), 2.38 (s, 3H), 2.30 (s, 3H), 1.65 (d, *J* = 6.7 Hz, 3H).

### Synthesis of Disulfanediylbis(ethane-1,1-diyl) bis(2-acetoxybenzoate) (HPD3)

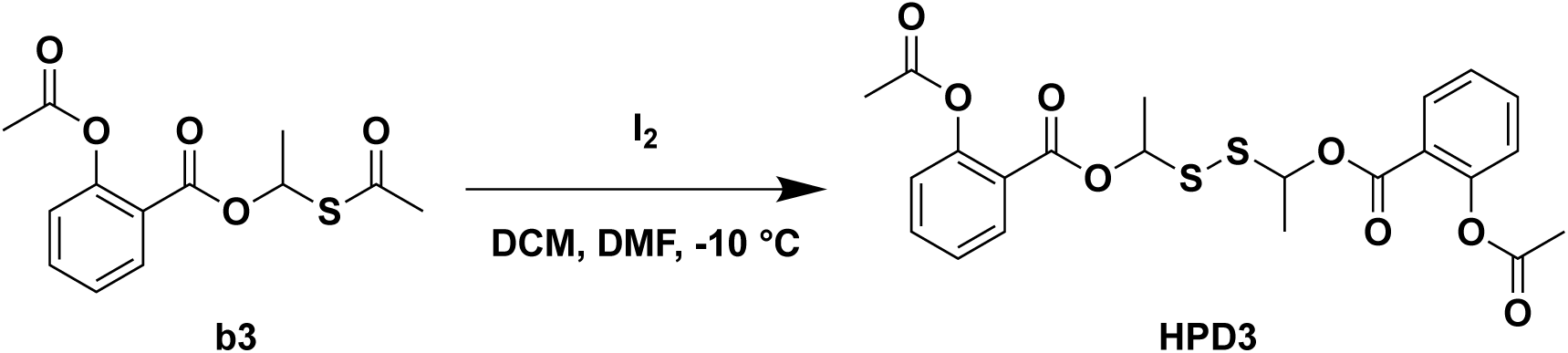

To a solution of 1-(acetylthio)ethyl 2-acetoxybenzoate (150 mg, 1.0 eq.) in DCM (2 mL) was added I_2_ (270 mg, 2.0 eq.) in DMF (1 mL) dropwise at-10 ℃. The mixture was then stirred at-10 ℃ for 1 hour and then warmed up to room temperature and stirred for 4 hours. The reaction was then monitored by TLC. After completion, the mixture was quenched with Na_2_S_2_O_3_ aq. and extracted with ethyl acetate for three times. The organic layer was separated and dried over Na_2_SO_4_ and concentrated in vacuo to afford a residue. The residue was purified by flash column chromatography (petroleum ether: ethyl acetate = 10: 1) to afford desired product as a colorless oil (80 mg, yield: 61.7%). ^1^H NMR (400 MHz, Chloroform-*d*) δ 8.02 – 7.92 (m, 2H), 7.55 – 7.45 (m, 2H), 7.28 – 7.21 (m, 2H), 7.08 – 7.00 (m, 2H), 6.18 – 6.08 (m, 2H), 2.28 (d, *J* = 7.9 Hz, 6H), 1.62 – 1.55 (m, 6H). ^13^C NMR (101 MHz, Chloroform-*d*) δ 169.64, 169.62, 163.22, 163.20, 150.89, 150.84, 134.30, 134.19, 132.01, 131.92, 126.14, 126.06, 123.96, 123.91, 122.77, 122.65, 78.16, 21.11, 21.08, 20.40, 20.32. (two signals observed for most carbons due to the presence of enantiomers). HRMS (ESI-QQQ) *m/z* [M + Na]^+^ Calculated for C_22_H_22_O_8_NaS_2_: 501.0648, found: 501.0624.

**Scheme S1.**
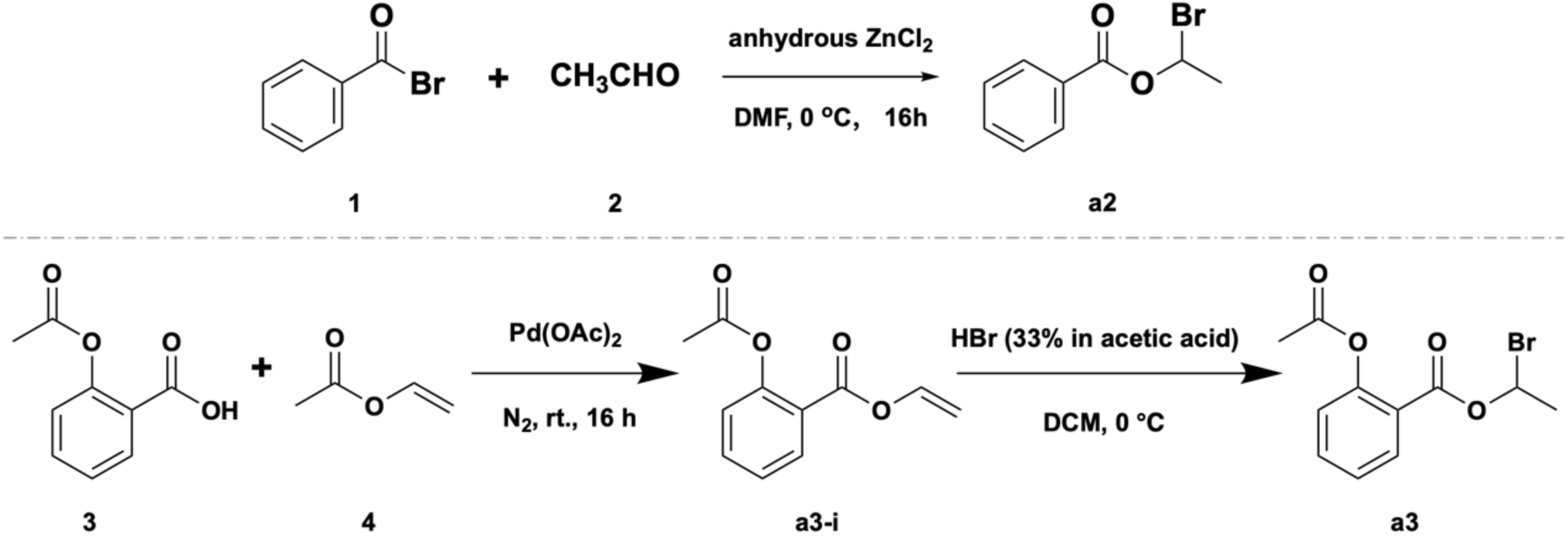
Synthetic Routes of starting materials a2-a3.

**Table S1.** Solubility in the Vehicle (DMSO:PEG400:H_2_O = 1:1:8) by HPLC.

| Comp. | M.W. | mg/mL |  | mM |  |
| --- | --- | --- | --- | --- | --- |
|  |  | Mean | SEM | Mean | SEM |
| **HPD1** | 238.32 | 1.8127 | 0.0538 | 7.6062 | 0.2259 |
| **HPD2** | 362.46 | 0.0037 | 0.0018 | 0.0102 | 0.0048 |
| **HPD3** | 478.53 | 0.0173 | 0.0008 | 0.0362 | 0.0017 |
| **HPS-2** | 558.6 | 0.0026 | 0.0003 | 0.0046 | 0.0006 |

\* st\_Area means standard solution dissolved in MeOH.

**Table S2.** Inhibition rate (%) of HPD1 at 10 μM against CYP2C9, CYP2D6, CYP3A4, CYP1A2, CYP2C19.

| Compound | CYP1A2 | CYP2C9 | CYP2C19 | CYP2D6 | CYP3A4<br>(Substrate:midazolam) | CYP3A4<br>(Substrate:testosterone) |
| --- | --- | --- | --- | --- | --- | --- |
| α-Naphthoflavone | 0.0219 | - | - | - | - | - |
| Sulfaphenazole | - | 0.304 | - | - | - | - |
| N-3-benzylnirvanol | - | - | 0.104 | - | - | - |
| Quinidine | - | - | - | 0.217 | - | - |
| Ketoconazole | - | - | - | - | 0.0287 | - |
|  | - | - | - | - | - | 0.0426 |
| **HPD1** |  |  |  |  |  |  |
| **Inhibition% at 10 µM** | **16.01** | **18.07** | **38.04** | **12.83** | **0.00** | **0.00** |

**Table S3.**
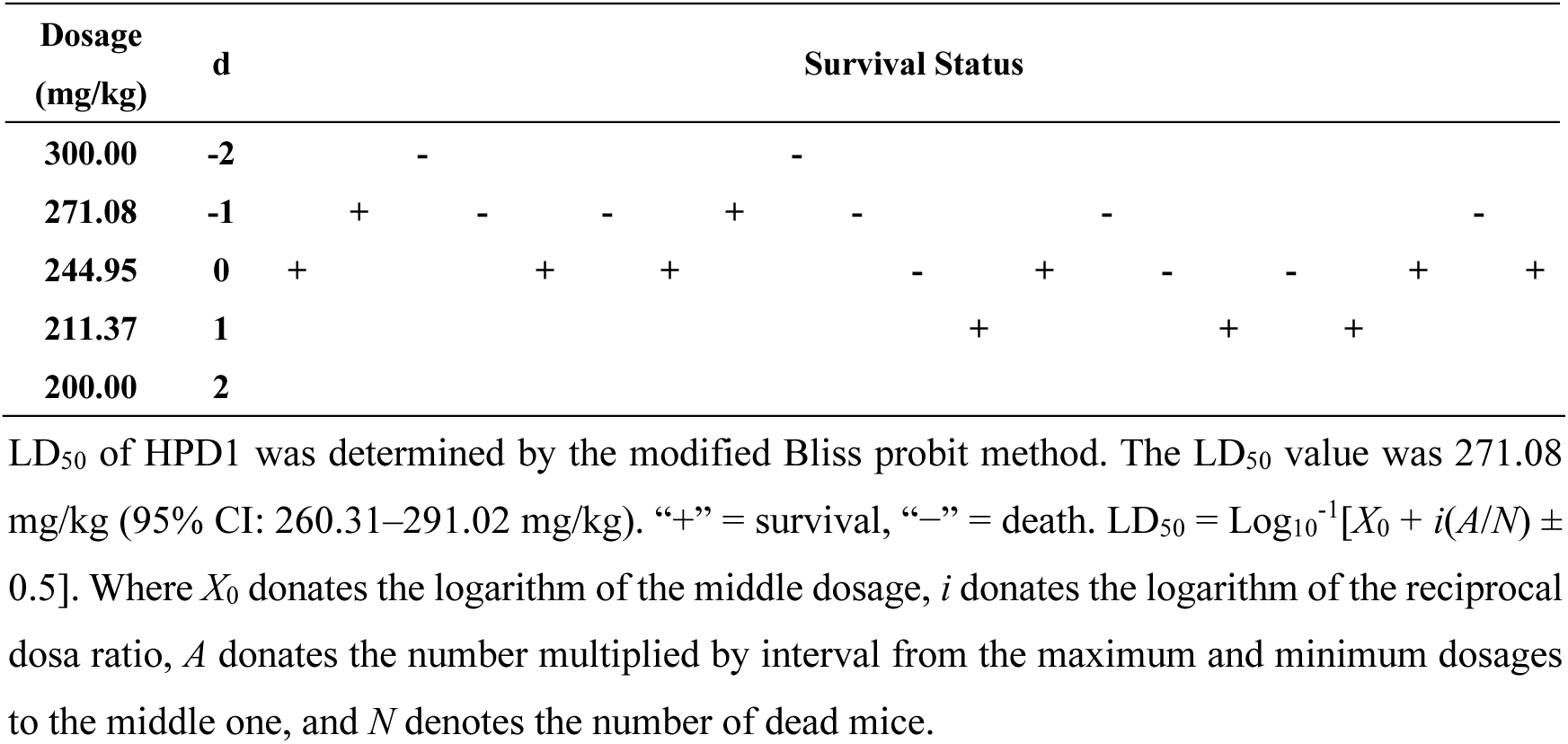
Survival and Mortality Data in LD_50_ Assay.

---

**Figure S1.**
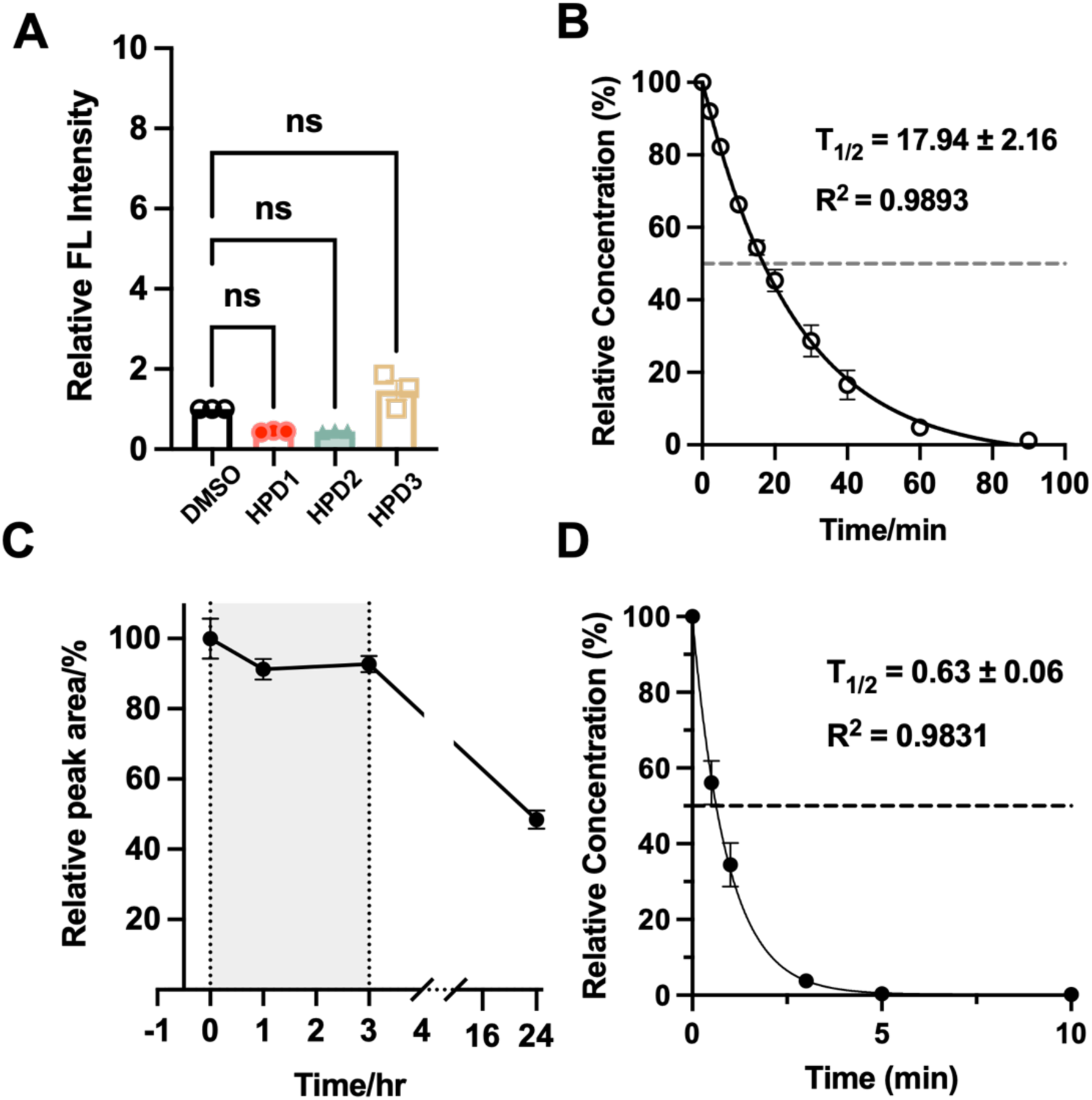
Characterization of Persulfide Precursors. (A) Semi-quantitative fluorescence analysis of H_2_S_2_ release from HPDs in the absence of PLE, using the DSP-3 probe λ_ex_/λ_em_ = 490/515 nm. Data are mean ± SEM (n = 3); ns, not significant (one-way ANOVA). (B) Pseudo-first-order kinetic fitting of HPD1 decomposition *in vitro*, yielding a half-life (T_1/2_) of 17.94 ± 2.16 min (one-phase decay model, R^2^ = 0.9893). (C) HPLC-based stability assessment of HPD1 in phosphate-buffered saline (PBS) over an extended period. (D) Kinetic analysis of HPD1 decomposition in mouse plasma, with a fitted half-life (T_1/2_)) of 0.63 ± 0.06 min (one-phase exponential decay model, R^2^ = 0.9831).

**Figure S2.**
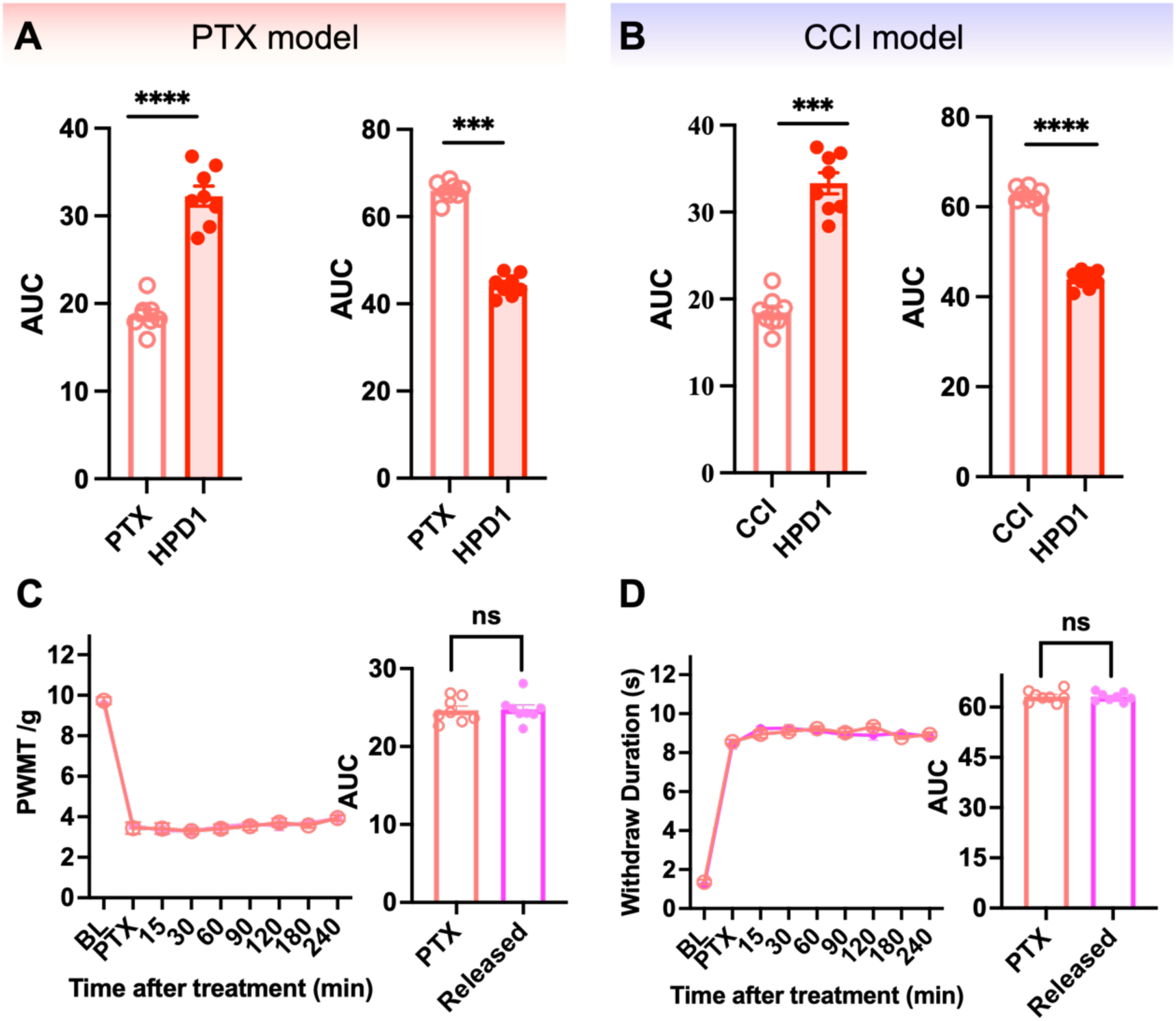
Behavioral assessment of HPD1 and its activation byproducts in neuropathic pain models. (A, B) Area under the curve (AUC) analysis of pain behavioral responses in the PTX (A) and CCI (B) models. Left panels: mechanical withdrawal thresholds (von Frey test); right panels: withdraw duration (acetone test). Data are presented as mean ± SEM (AUC values). (C, D) Time-course (left) and AUC (right) analyses of mechanical (C) and cold (D) pain thresholds in the PTX model. Mice received either vehicle or the HPD1-released mixture. Statistical analyses were performed using two-tailed unpaired Student’s t-test. ns, not significant; ***p < 0.001, ****p < 0.0001.

**Figure S3.**
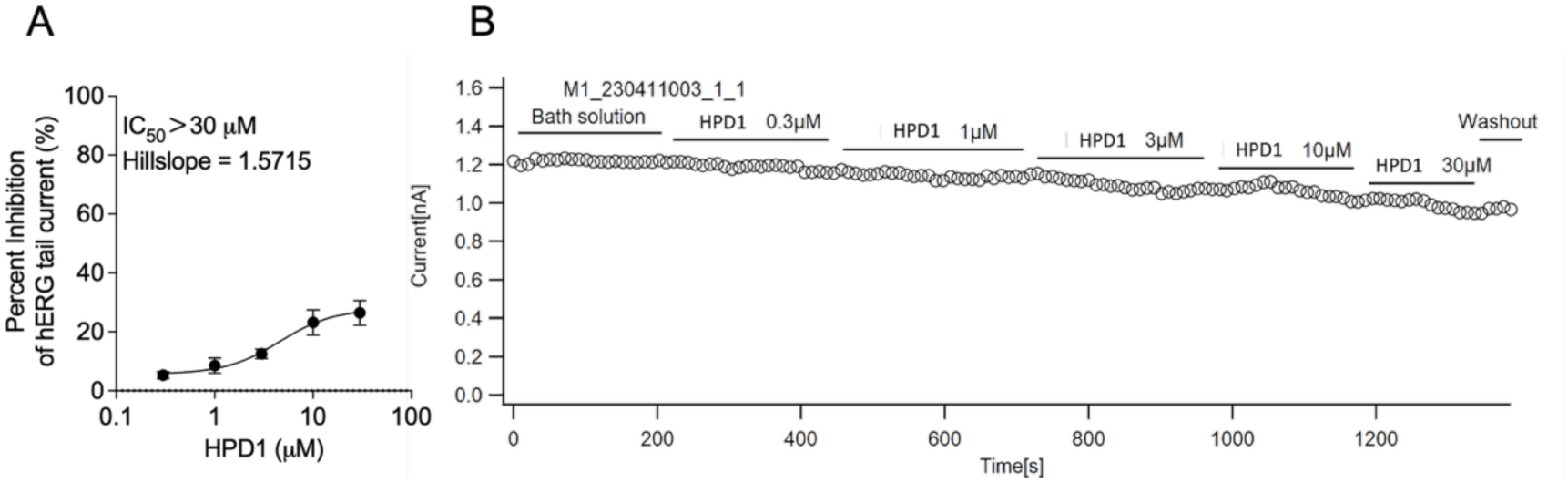
Inhibitory effects of HPD1 on hERG channel activity. (A) Concentration-response curve for HPD1-mediated inhibition of hERG tail current. Data are expressed as mean ± SEM (n = 3). The half-maximal inhibitory concentration (IC_50_) was determined to be > 30 μM, with a Hill slope of 1.5715. (B) Representative time course of hERG current recorded from a single cell during sequential perfusion of bath solution and increasing concentrations of HPD1 (0.3, 1, 3, 10, and 30 μM), followed by washout. Current amplitude was normalized to the baseline value recorded in bath solution.

**Figure S4.**
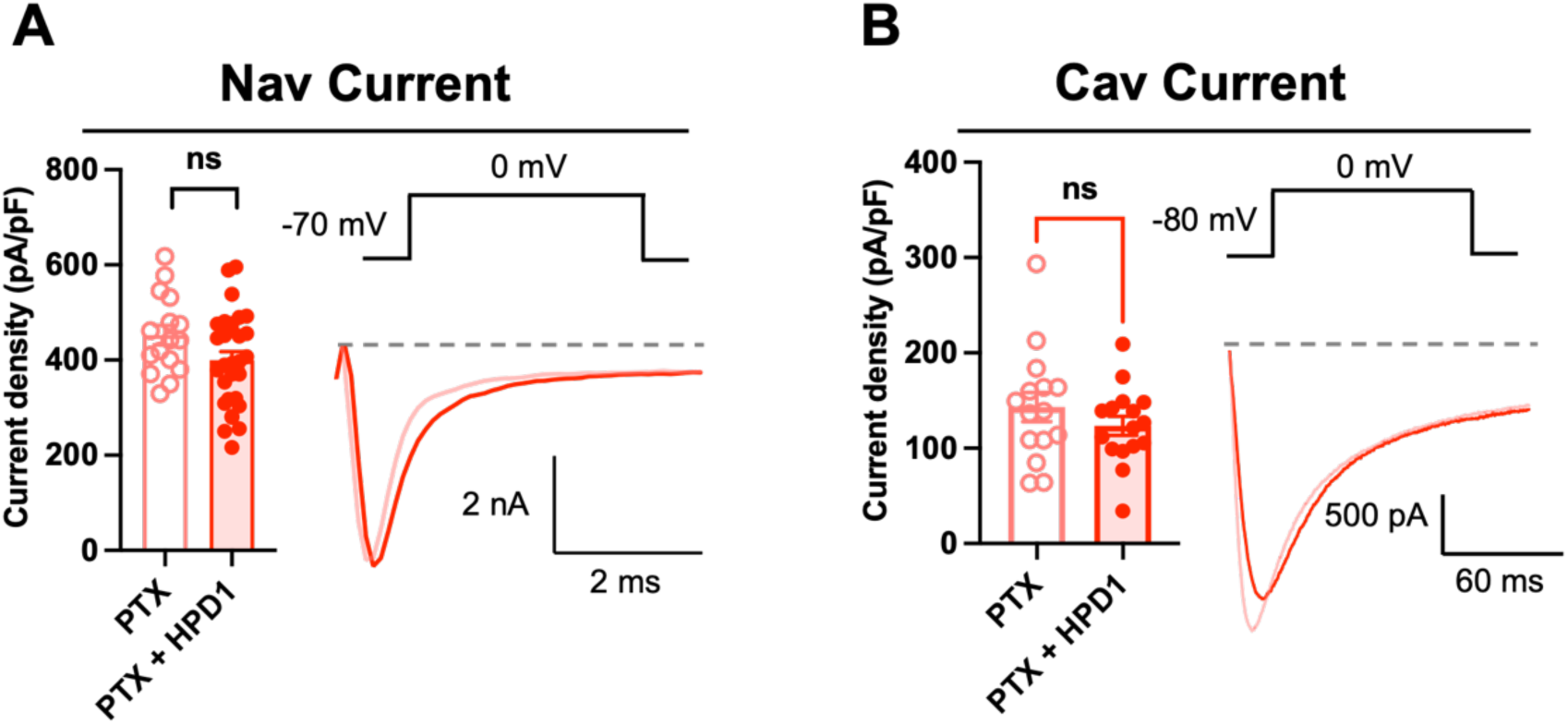
Effects of HPD1 on voltage-gated currents in DRG neurons. (A) Voltage-gated sodium (Nav) currents, evoked from-70 mV holding potential to 0 mV for 50 ms, with summary current density and representative traces. (B) Voltage-gated calcium (Cav) currents, evoked from - 80 mV holding potential to 0 mV for 200 ms, with summary current density and representative traces. Statistical analyses were performed using two-tailed unpaired Student’s t-test. ns, not significant.

**Figure S5.**
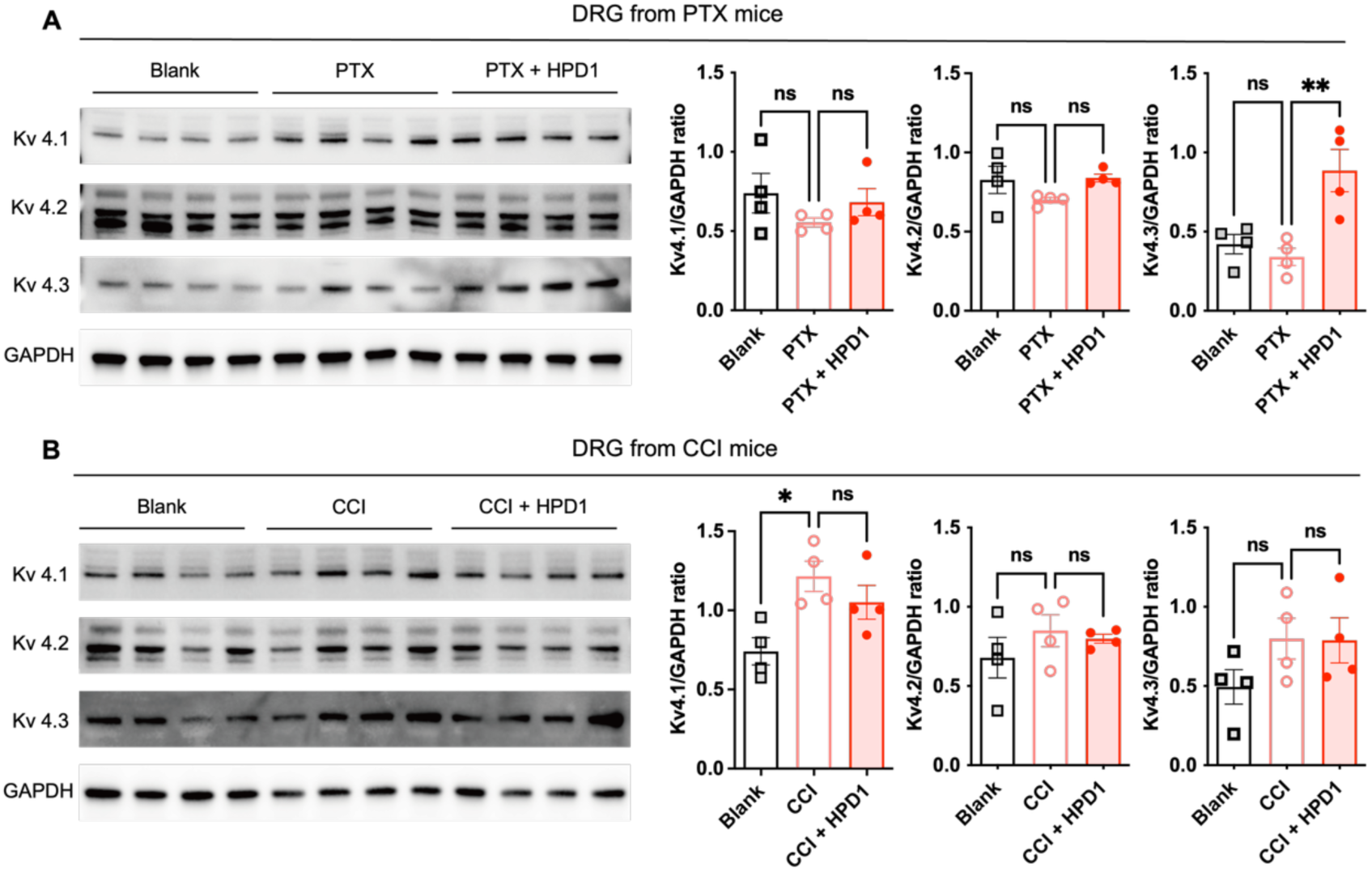
Western blot analysis of Kv4.1, Kv4.2, and Kv4.3 protein expression in DRG tissues. (A, B) Representative immunoblots (left) and corresponding quantitative analysis (right) in mice with PTX-induced (A) and CCI-induced (B) neuropathic pain, following the indicated treatments. Band intensities were normalized to GAPDH. Data are mean ± SEM (n = 4). One-way ANOVA followed by Tukey’s multiple comparisons test. ns, not significant; *p < 0.05; **p < 0.01.

**Figure S6.**
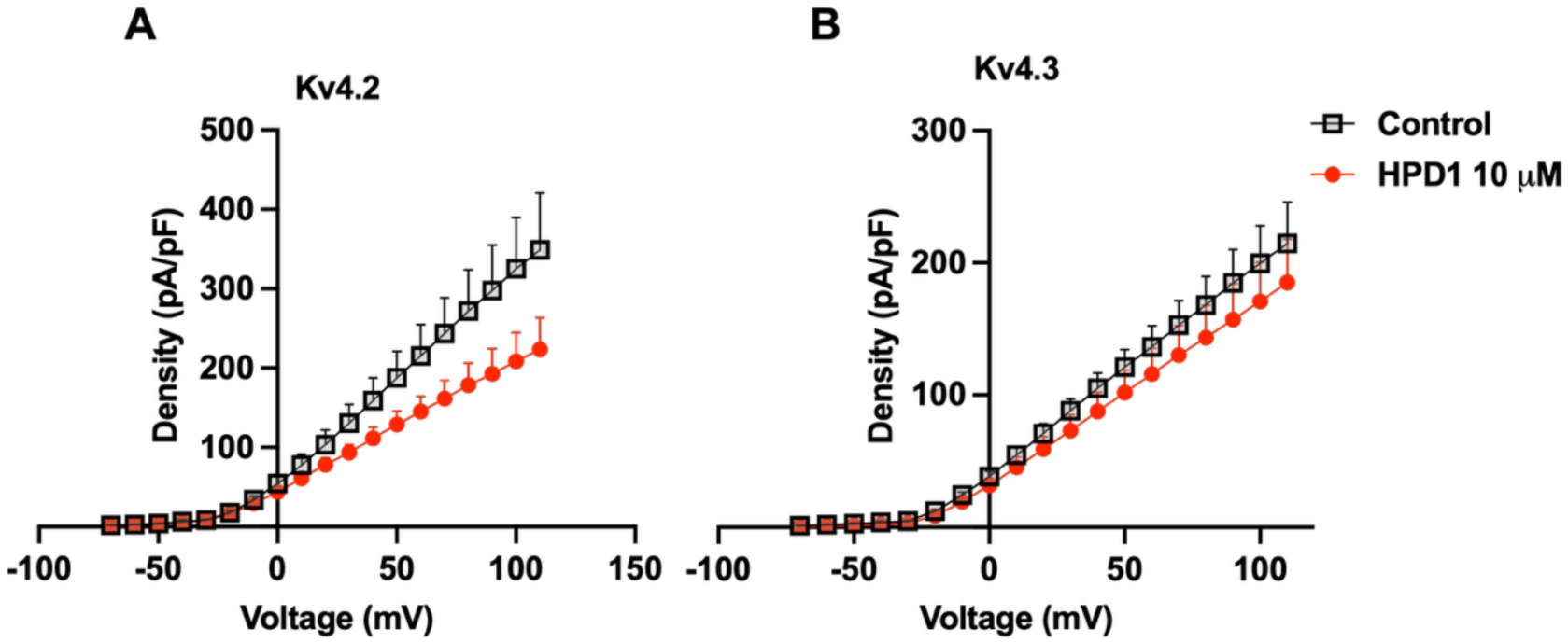
Effect of HPD1 on recombinant Kv4 subtype currents. I–V curves of Kv4.2 (A) and Kv4.3 (B) channels stably expressed in HEK293 cells, recorded under control conditions and after bath application of 10 μM HPD1. Current density (pA/pF) is whole-cell current normalized to membrane capacitance. Data are mean ± SEM (n ≥ 3 cells per group).

**Figure S7.**
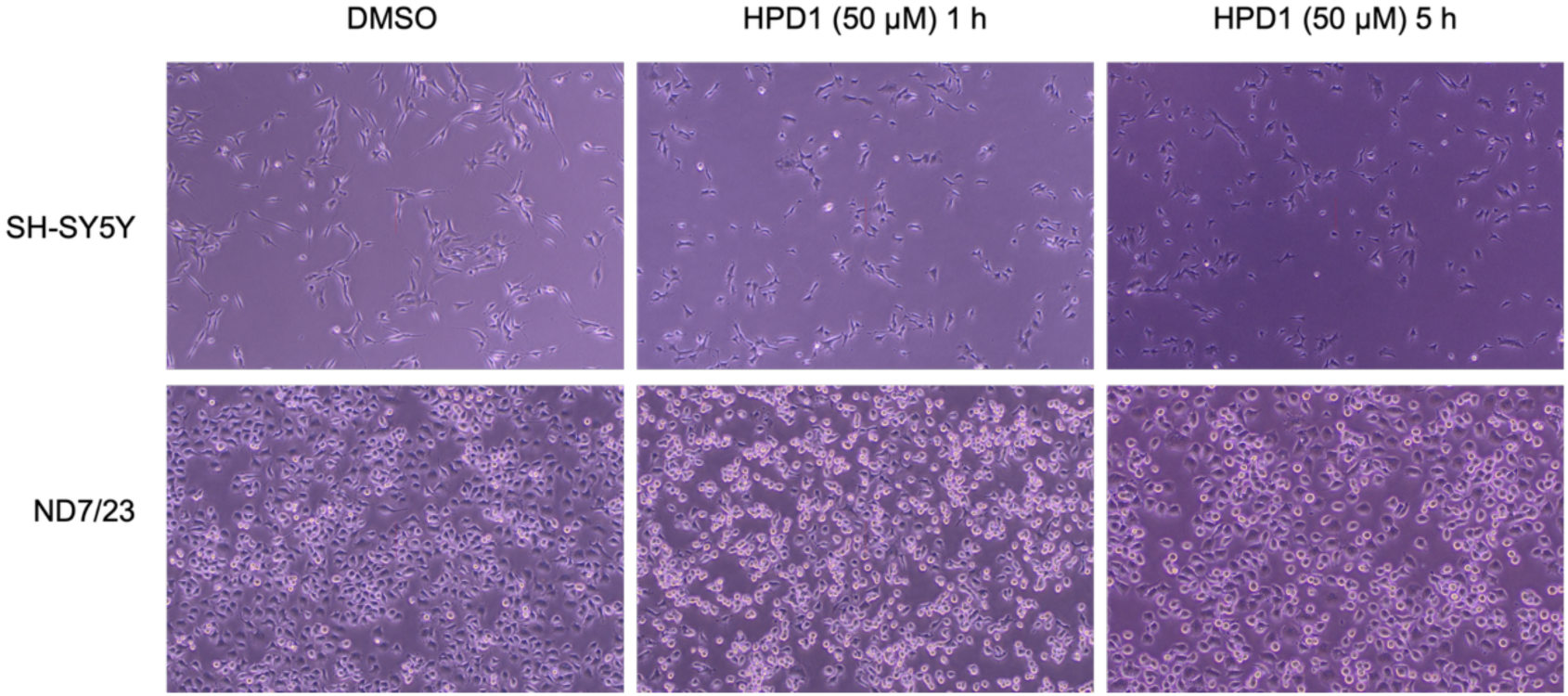
Morphology of SH-SY5Y and ND7/23 cells after HPD1 treatment. Cells treated with 50 μM HPD1 or DMSO for the indicated time points observed under a light microscope (10 × objective).

**Figure S8.**
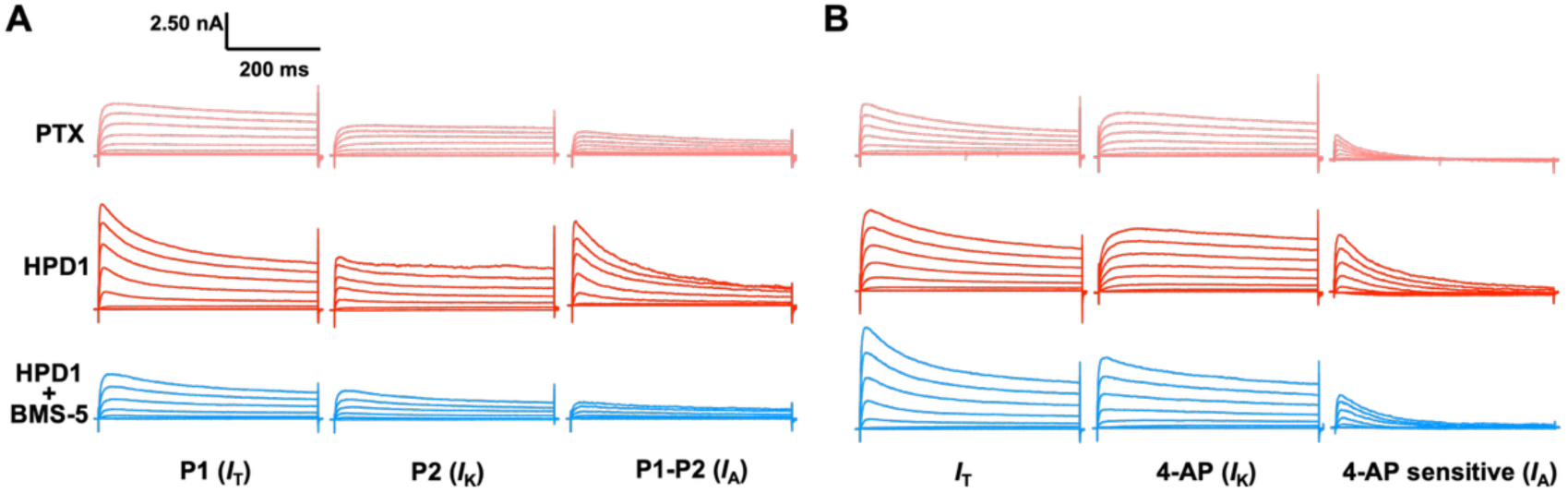
Representative traces of isolated A-type potassium currents (*I*_A_). (*I*_A_) isolated from DRG neurons using two different protocols. (A) *I*_A_ isolation via a two-step voltage protocol. (B) *I*_A_ isolation via pharmacological subtraction with 4-aminopyridine (4-AP). For both protocols, currents are shown under three conditions: PTX control, HPD1 (10 μM) treatment, and HPD1 (10 μM) + BMS-5 (1 μM) co-treatment. Black traces denote total current (*I*_T_), gray traces denote sustained current (*I*_K_), and red traces represent isolated *I*_A_. Scale bars: 2.50 nA, 200 ms.

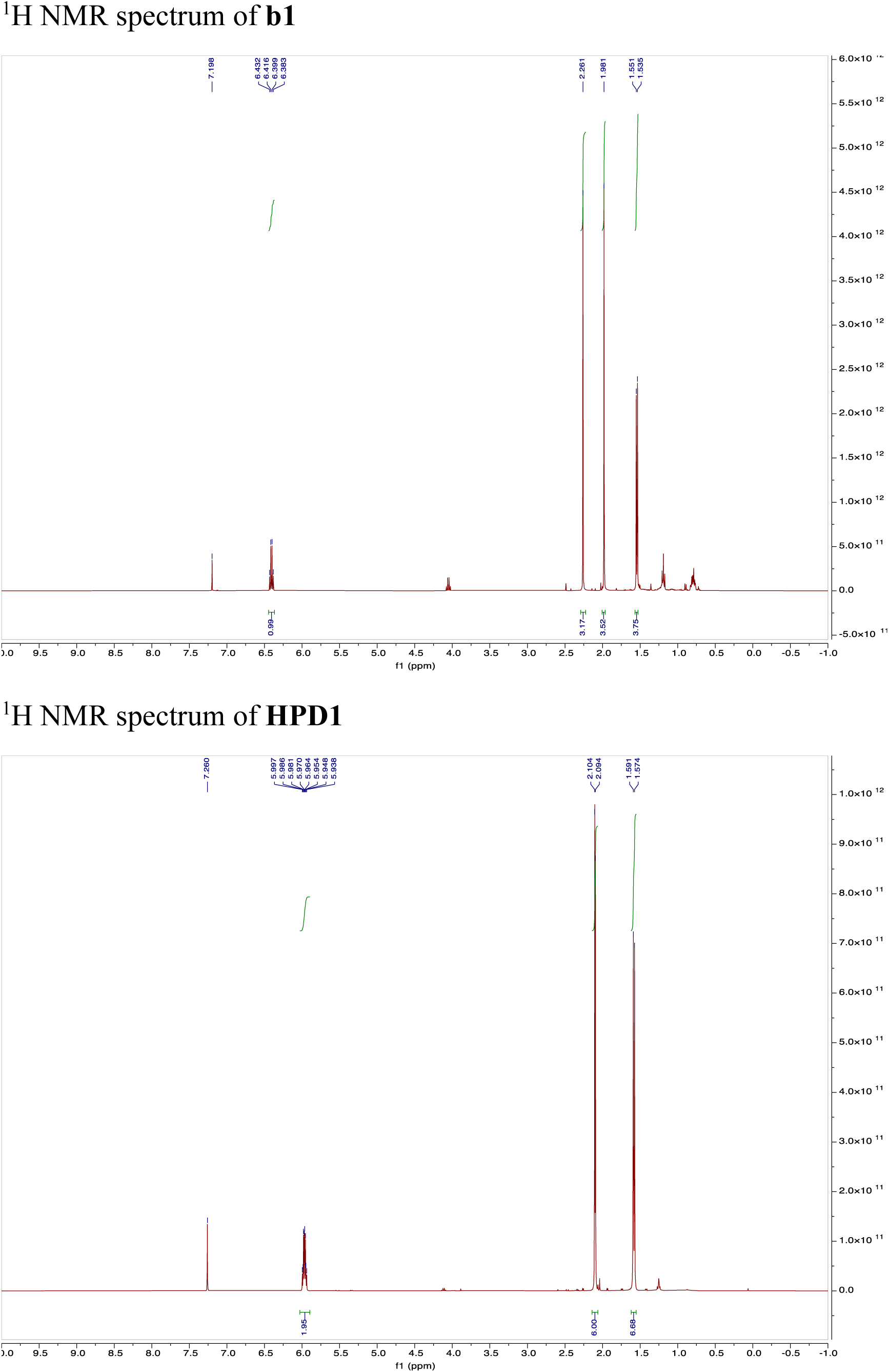

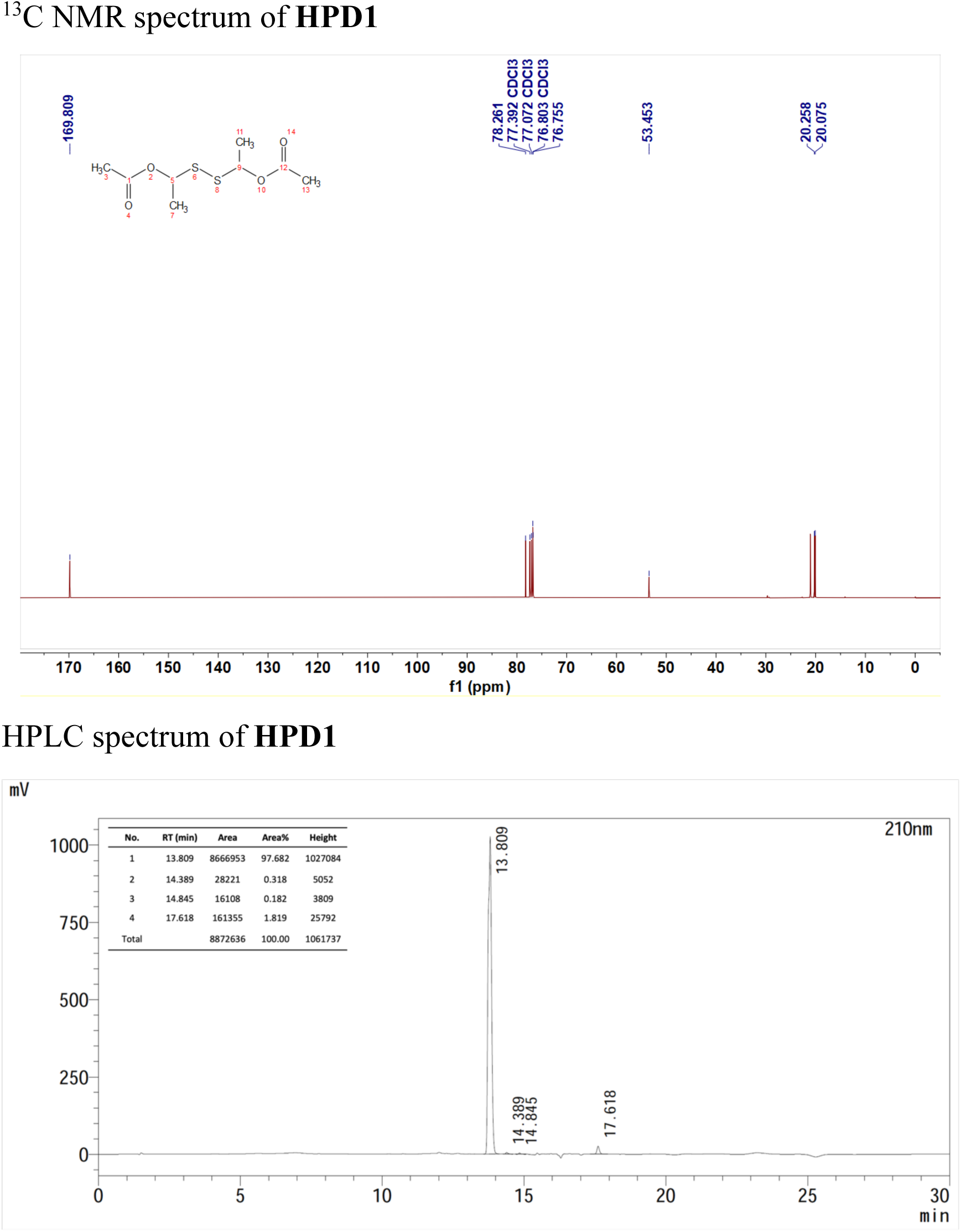

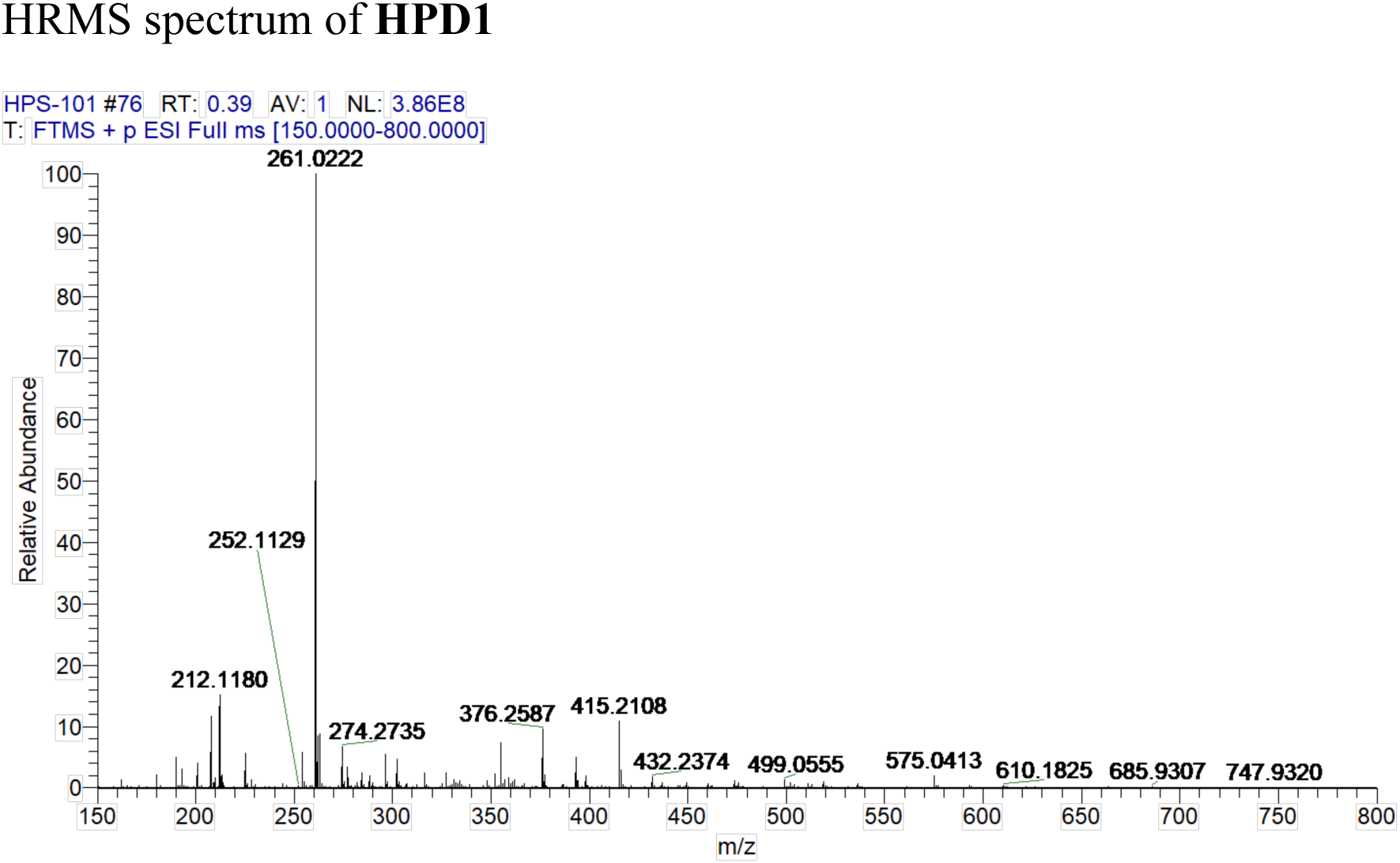

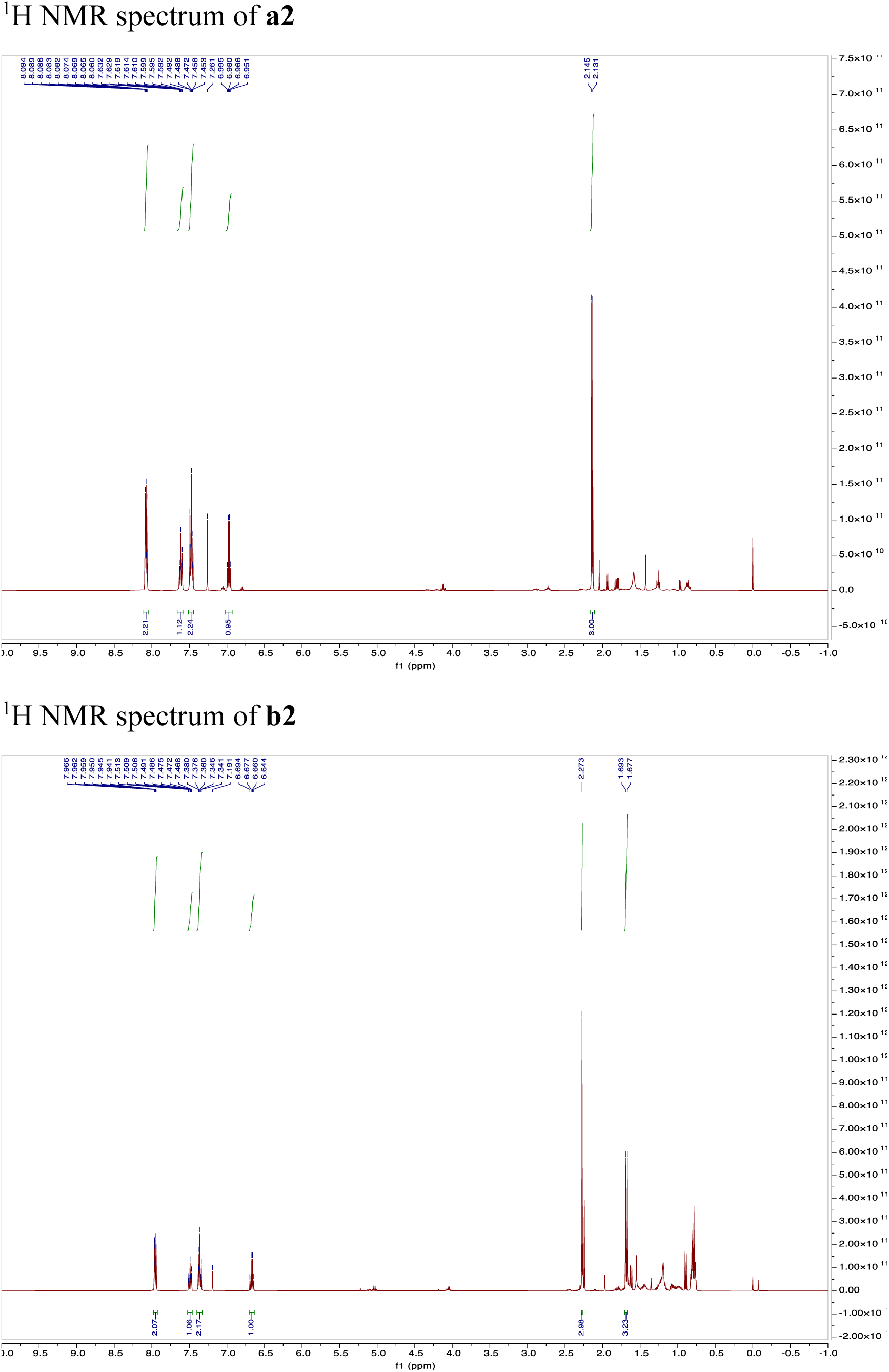

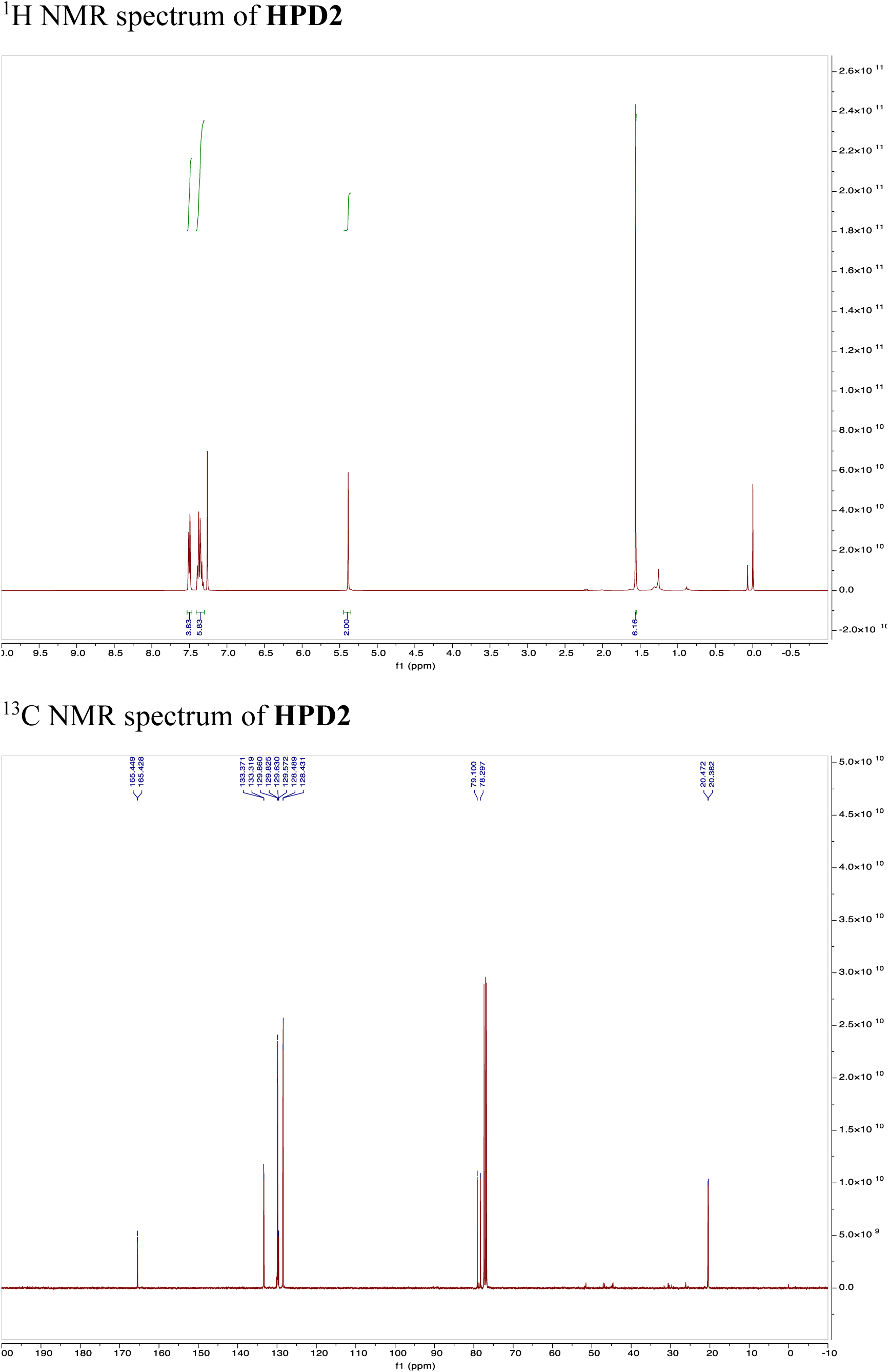

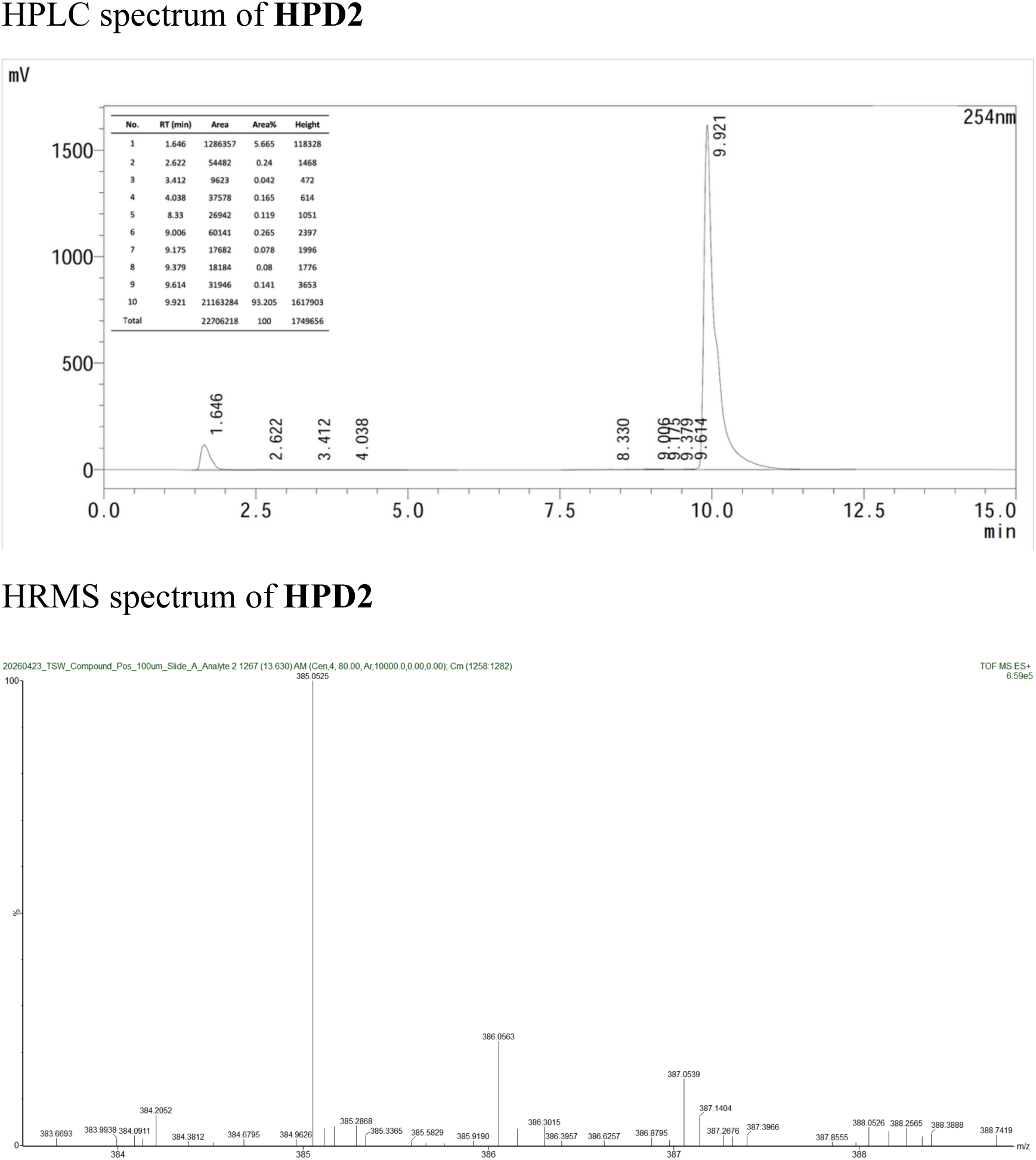

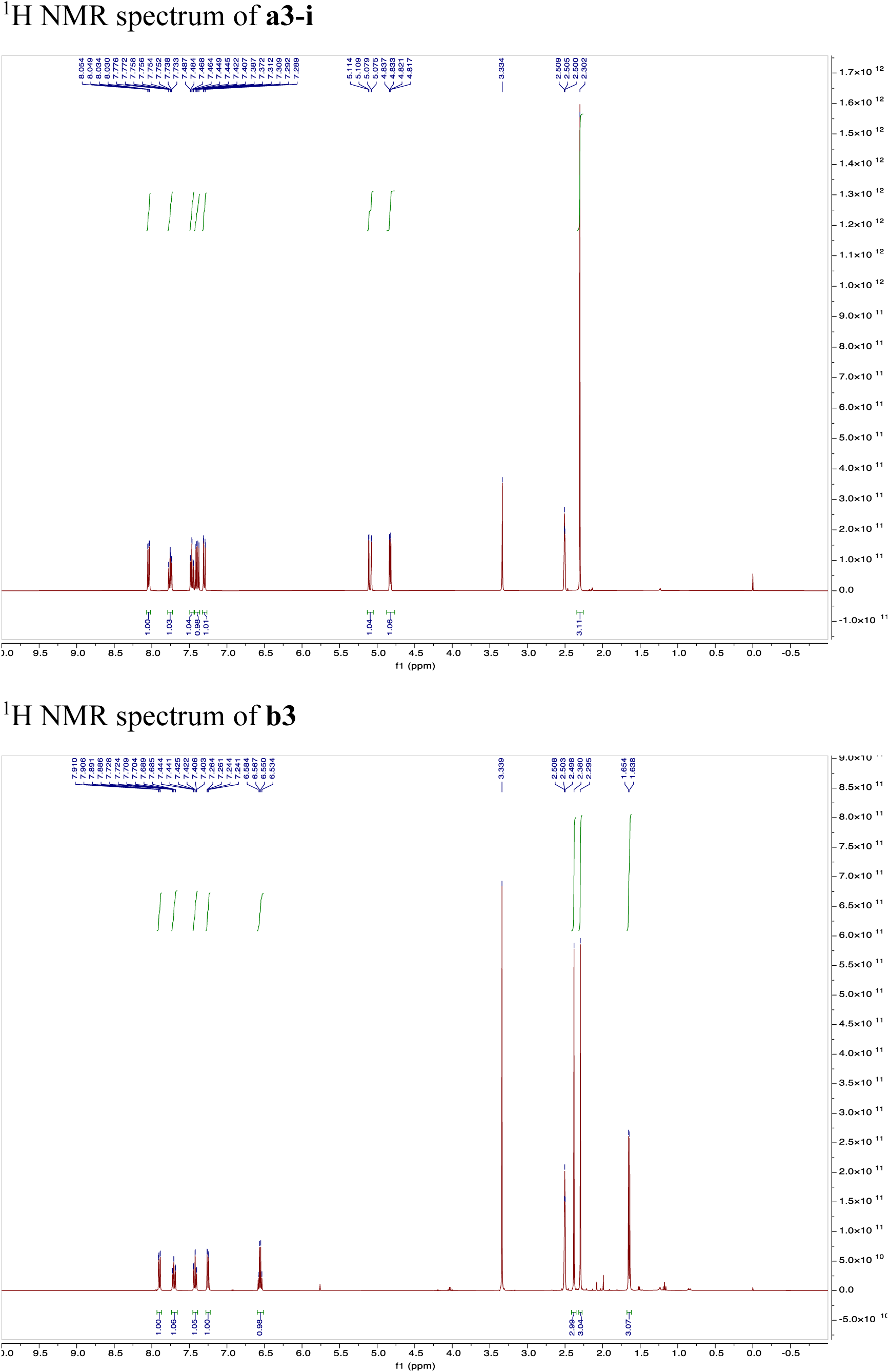

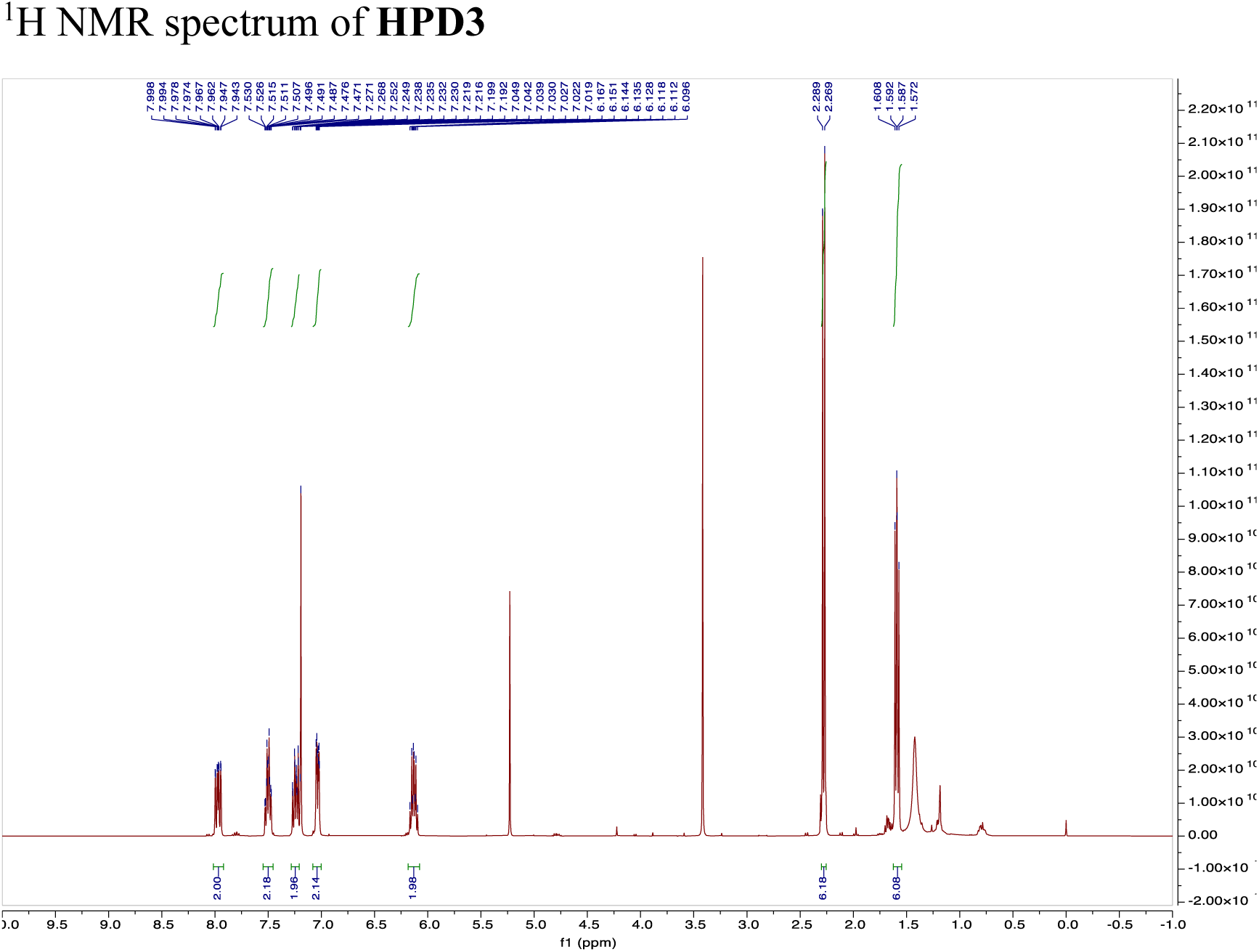

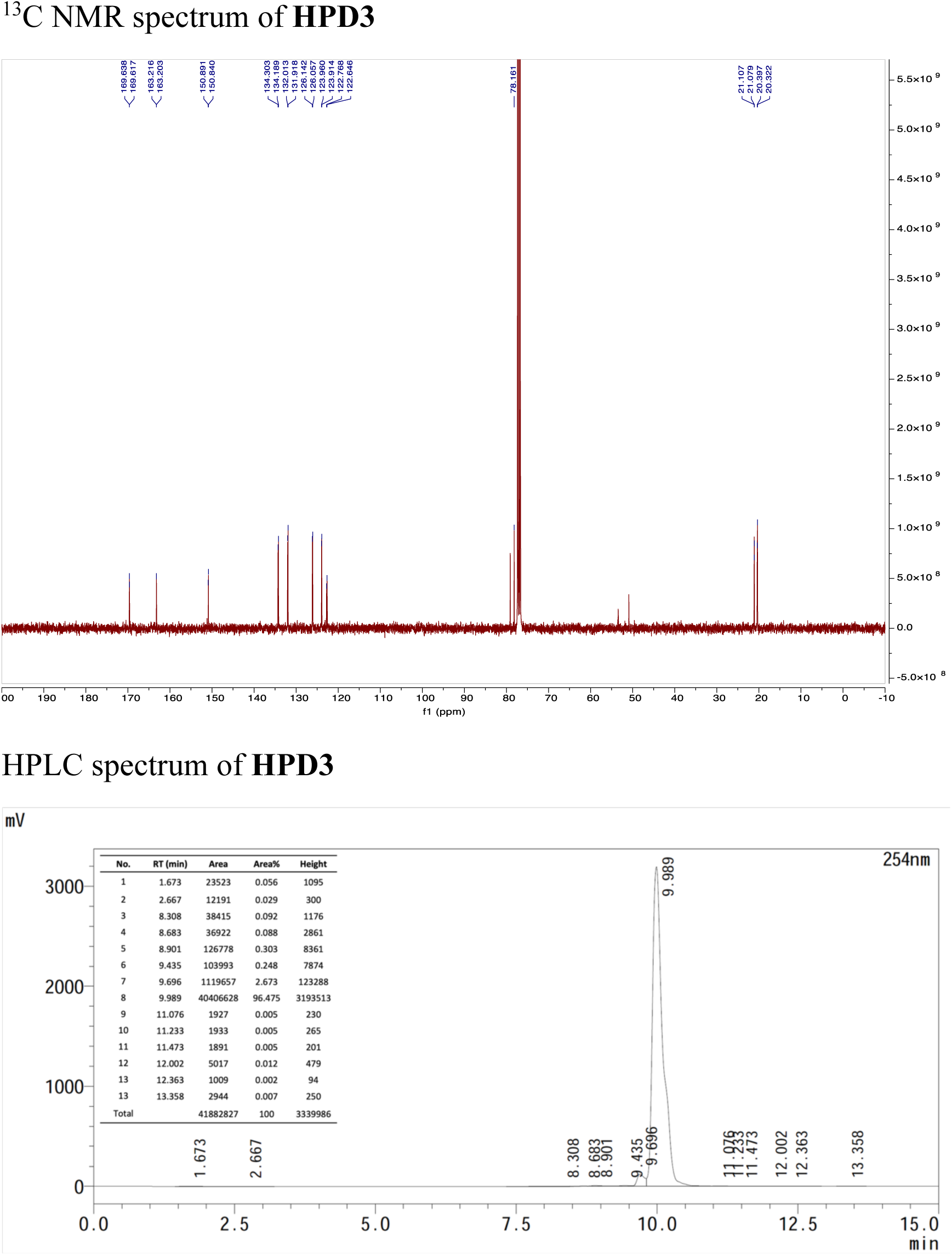

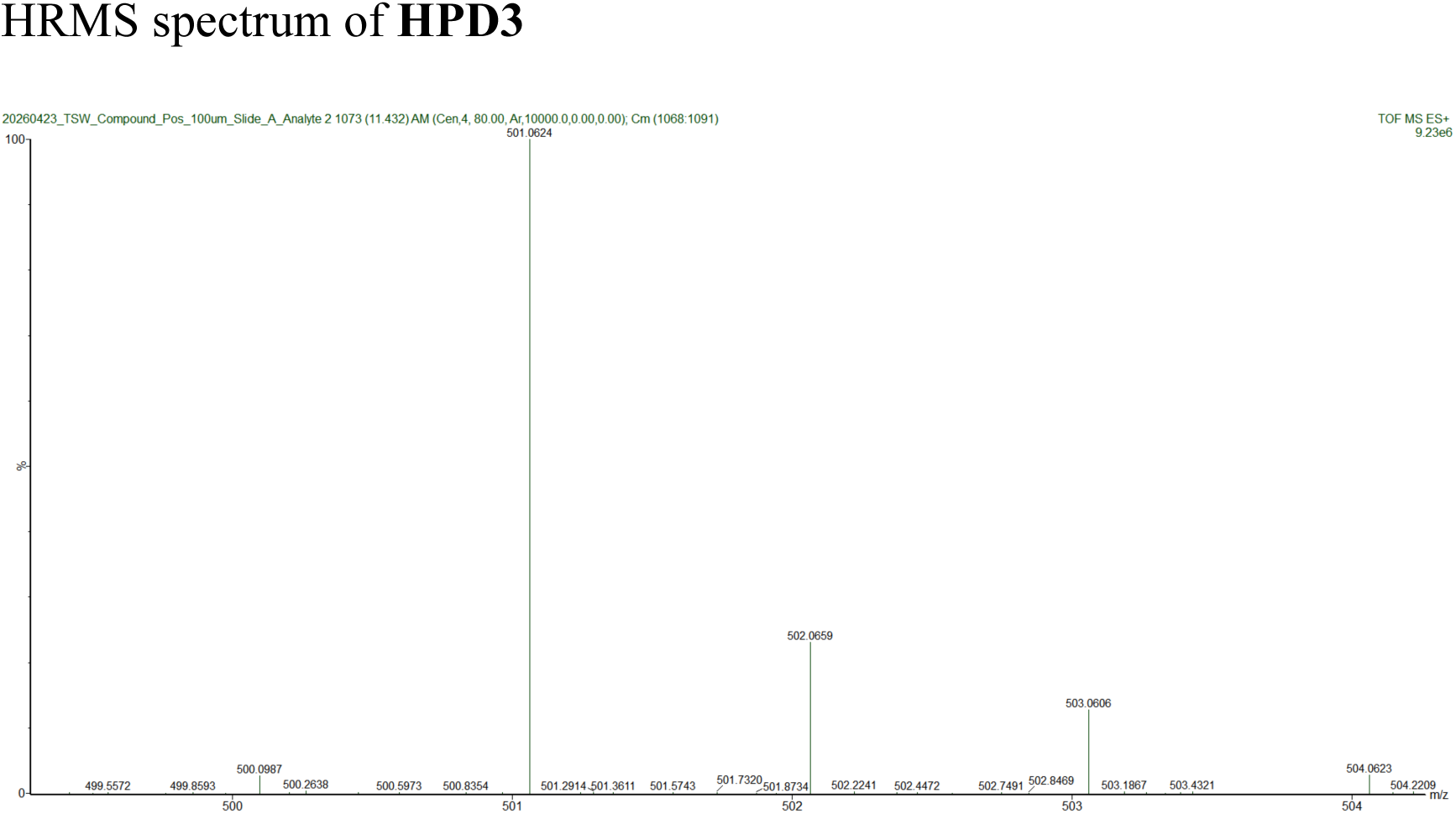

